# Chiral monoterpene dynamics of shoots and roots of Norway spruce in response to drought

**DOI:** 10.1101/2025.05.02.651829

**Authors:** L. Erik Daber, Jürgen Kreuzwieser, Mirjam Meischner, Jonathan Williams, Christiane Werner

## Abstract

Although chiral monoterpenes emitted by plants above- and belowground shape the chemical landscape of many ecosystems, their biosynthesis and emissions, especially in response to drought, are poorly understood. We imposed a 6-week drought on two-year old, potted saplings of Norway spruce and analysed chiral monoterpene emissions and tissue concentrations from needles and roots. Isotopically labelled pyruvate was used to compare tissue-specific contributions of *de novo* synthesis to chiral monoterpene concentrations. While *de novo* synthesis of (−)-α-pinene and both enantiomers of limonene was apparent in needle emissions, no label was incorporated in roots. Drought reduced chiral monoterpene emissions to 30% of control levels, but increased needle and root tissue concentrations by 150 and 230%, respectively. Aboveground monoterpene concentrations were dominated by (−)-limonene, whereas belowground concentrations mainly consisted of the (−)-enantiomers of α-pinene, β-pinene, β-phellandrene and camphene. Chiral composition in needles shifted in response to drought but remained stable in roots. We conclude that chiral monoterpene composition is tissue-specific and likely related to tissue-specific functioning. Instead of being passively emitted from storage pools, our results suggest active control mechanisms regulating chiral monoterpene emissions under drought conditions. Our findings imply important ramifications for understanding the regulation of emissions in relation to storage pools and plant-environmental interactions.

**Summary Statement:** Emissions of chiral monoterpenes and their composition is tissue-specific and regulated independent from storage pools in *Picea abies*. Chiral monoterpene ratios shift aboveground in response to drought, but are not affected belowground.

## Introduction

Plants emit a plethora of monoterpenes, many of which exist in mirror images, also called enantiomers. These chiral forms of monoterpenes play distinct roles in plant-insect interactions (de los Santos and Wolf, 2020; Phillips, Savage and Croteau, 1999; Rufino *et al.*, 2014) and their enantiomeric ratios can be utilised to detect plant physiologic stress responses such as drought (Byron *et al.*, 2022) and mechanical wounding (Williams *et al.*, 2011; Yassaa and Williams, 2007) in various ecosystems and plant genera. Much attention in early works has been given to plant species with specialised storage structures, such as resin ducts; due to their high concentration and diversity of chiral monoterpenes (Borg-Karlson *et al.*, 1993; Norin, 1996; Persson, Borg-Karlson and Norin, 1993). Even though differences in enantiomeric ratios of monoterpenes in storage pools of shoots and roots of conifer species have been studied (Persson, Borg-Karlson and Norin, 1993; Sjödin *et al.*, 1996), the relation of chiral monoterpene emissions to their storage pools and biosynthesis in response to drought, remains unresolved. Biological activity and antimicrobial and/ or insecticidal properties of chiral monoterpenes can differ between enantiomers (Aggarwal *et al.*, 2002; Cheng *et al.*, 2017; Da Rivas Silva *et al.*, 2012; Hachlafi *et al.*, 2023) and might be enhanced depending on enantiomeric ratios (Neirotti, Moscatelli and Tiscornia, 1996; van Vuuren and Viljoen, 2007). Given that drought might affect chiral ratios of monoterpene emissions and storage pools, possible shifts in chiral ratios could be utilized by pest species, such as bark beetles, as cues to identify weakened tree individuals during primary attraction (Lehmanski *et al.*, 2023).

Chiral monoterpenes are synthesized within the plastid by monoterpene synthases (mTPS), many of which have been identified over the last decades (Aubourg, Lecharny and Bohlmann, 2002; Chen *et al.*, 2011; Keeling *et al.*, 2011; Zhou and Pichersky, 2020). In the case of Norway spruce, four multi-product- and one single-product mTPS were identified and their activity within foliage and stem has been affirmed (Fäldt *et al.*, 2003; Martin, Fäldt and Bohlmann, 2004). Norway spruce (Borg-Karlson *et al.*, 1993; Duan et al., 2020) and many other plant species also accumulate monoterpenes in their roots (Bos *et al.*, 2002; Wichtmann and Stahl-Biskup, 1987; Kleiber et al., 2017; Duan et al. 2019) and partially emit them into the rhizosphere (Janson, 1993; Lin, Owen and Peñuelas, 2007; Steeghs *et al.*, 2004), where they play important roles in deterring herbivores and pathogens (Bernard-Dagan, 1988; Krupa and Fries, 1971; Stotzky and Schenck, 1976; Unsicker, Kunert and Gershenzon, 2009), and might be utilised as allelopathic agents (Chen *et al.*, 2004; Delory *et al.*, 2016). However, enantiomeric ratios of root-released monoterpenes have not been investigated until now. Roots produce a diverse spectrum of secondary metabolites, but depend on a supply of photosynthates from the shoot (Flores, Vivanco and Loyola-Vargas, 1999). Several active mTPS have been characterised in roots of different plant species (Chen *et al.*, 2004; Zhou and Pichersky, 2020), indicating shoot-independent biosynthesis of monoterpenes in roots. Biosynthesis is most likely located in root plastids driven by the MEP-pathway (Hans *et al.*, 2004); more precisely in leucoplasts of secretory cells of resin ducts in conifer species (Cheniclet and Carde, 1985; Schürmann *et al.*, 1993). However, biosynthesis of monoterpenes might also depend on isoprenoid precursors being translocated via the phloem from the shoot towards roots (Burlat *et al.*, 2004).

Biosynthesis, storage and emission of chiral monoterpenes are affected by plant physiological, and physicochemical controls (Niinemets, Loreto and Reichstein, 2004). In species without specialised storage structures (such as resin ducts) monoterpene emissions are primarily limited by *de novo* synthesis, but additionally rely on non-specific storage pools in the lipid phase of the leaf, that can sustain emissions for 10-15 min without *de novo* synthesis (Loreto *et al.*, 1998; Niinemets and Reichstein, 2002). Terpenoid emission from species with specialised storage compartments were initially seen to be passively released from these storage compartments, only controlled by compound volatility (Guenther *et al.*, 1993), which is strongly affected by temperature (Fischbach *et al.*, 2002). However, several studies show that the suite of monoterpenes emitted by e.g. conifer species is partially light dependent and emitted *de novo*, and therefore only partially temperature dependent and emitted from storage compartments (Loreto *et al.*, 2000; Shao *et al.*, 2001; Staudt *et al.*, 1997). The contribution of *de novo* synthesis to total monoterpene emissions differs between conifer species (*Picea abies*: 33.5%; *Pinus sylvestris*: 58%; *Larix decidua*: 9.8% (Ghirardo *et al.*, 2010)) and depends on environmental conditions (Niinemets, Loreto and Reichstein, 2004) and stressors, diminishing under long-term drought stress (Lewinsohn *et al.*, 1993; Trapp and Croteau, 2001).

To elucidate how above- and belowground biosynthesis, emission and storage of chiral monoterpenes is affected by drought, we conducted a drought experiment on two-year old, potted saplings of Norway spruce (*Picea abies* L.) and analysed chiral monoterpene emissions and tissue concentrations of needles and roots. To estimate the contribution of monoterpene *de novo* synthesis to the overall emission and storage of needles and roots, we conducted stable C isotope tracer experiments with the central metabolite pyruvate. In particular, we were interested to investigate if and how chiral mixtures of monoterpenes emitted and stored by needles and roots differ, and how compositions are affected by drought. In the face of drought, the plant will adjust internal cycling of carbon resources and chiral changes can reveal how these processes change. If the contribution of *de novo* synthesis to total monoterpene emissions diminishes with drought, remaining emissions and their chiral composition should correspond with tissue concentrations. We therefore hypothesise that (*i*) contribution and chiral composition of stored monoterpenes to overall monoterpene emissions is determined by the monoterpene content of the tissue. Regarding belowground dynamics, we hypothesise that (*ii*) monoterpene emissions from roots are partially derived from *de novo* synthesis, but will be driven by monoterpene concentrations of storage compartments.

## Materials and Methods

### Plant material and drought treatment

Two-year old spruce-saplings (50-60cm) from a forest nursery (Burger, Zell am Harmersbach, Germany) were potted (5.6 L, 3:2 peat and quartz sand, 27.5 g NPK fertiliser) in a glasshouse and well-watered three times per week. Two weeks before the experiment they were transferred to climate chambers (16h/8 h, light/dark, 600-750 μmol m^-2^ s^-1^ (photosynthetically active radiation), T_day_: 22 °C, T_night_: 18°C) for acclimatisation. 24 plants were regularly irrigated to a volumetric water content (VWC) of ∼0.3 m^3^m^-3^ two weeks before the start of the experiment. Afterwards, plants were randomly split into control (VWC of ∼0.3 m^3^m^-3^ throughout the experiment) and drought (consistent reduction of VWC from ∼0.3 to ∼0.075 m^3^m^-3^within 21 days) treatments (for measurement schedule, see **Fig. 1**). Every other day, plants were weighed and watered accordingly to their group. VWC was measured at three different locations of each pot at 0-10cm depth, using a soil moisture sensor (Theta Probe ML3, delta T, Cambridge, England) (for details, see Daber *et al*. 2023, *in review*).

**Fig. 1.**
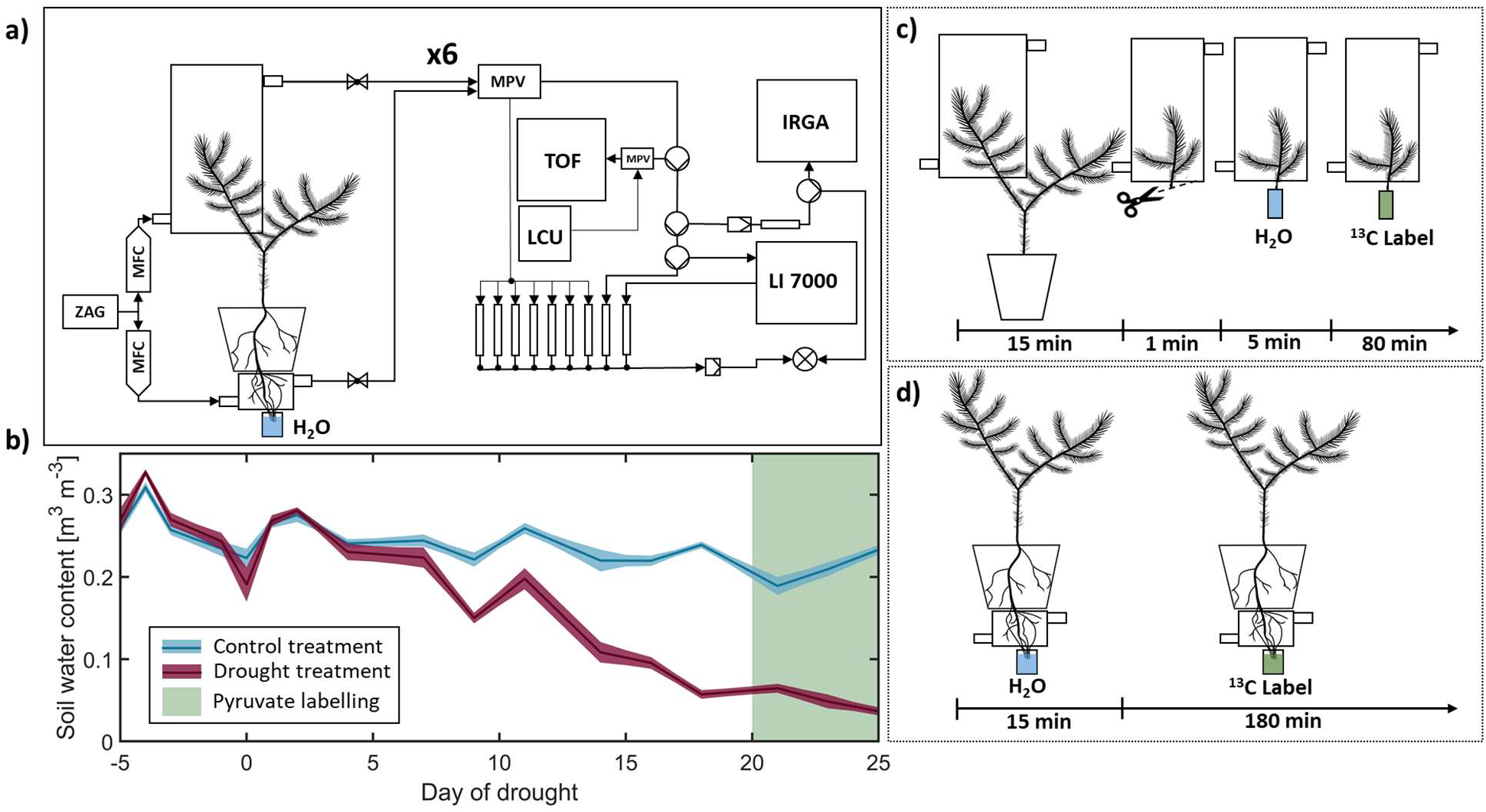
Schematic of the experimental design of the drought experiment with saplings of *Picea abies*. a) Setup inside of a walk-in climate chamber for parallel online measurements on 6 branches and roots of Norway spruce that were installed in flow-through chambers. Emissions were measured using a proton-transfer-reaction time-of-flight mass spectrometer (TOF), an isotopic gas ratio analyzer for CO_2_ (IRGA) and water vapour H_2_O (LI 7000). LCU, liquid calibration unit; MFC, mass flow controller; MPV, multi-position valve; ZAG, zero-air-generator (see Fasbender *et al.*, 2018). b) Soil water content of drought impacted (n =12, purple) and control (n = 12, blue) plants throughout the drought experiment. c) branch pyruvate labelling performed after 20 days of drought. After control measurements, installed branches were cut, placed for 5 minutes in distilled water, and afterwards placed in 10 mM ^13^C2-pyruvate solution. d) Root pyruvate labelling performed after 20 days of drought. After excavation, roots were placed in glass chambers, with the root tips being placed in H_2_O. Water was exchanged with 80 mM ^13^C2-pyruvate solution after control measurements.

**Fig. 2.**
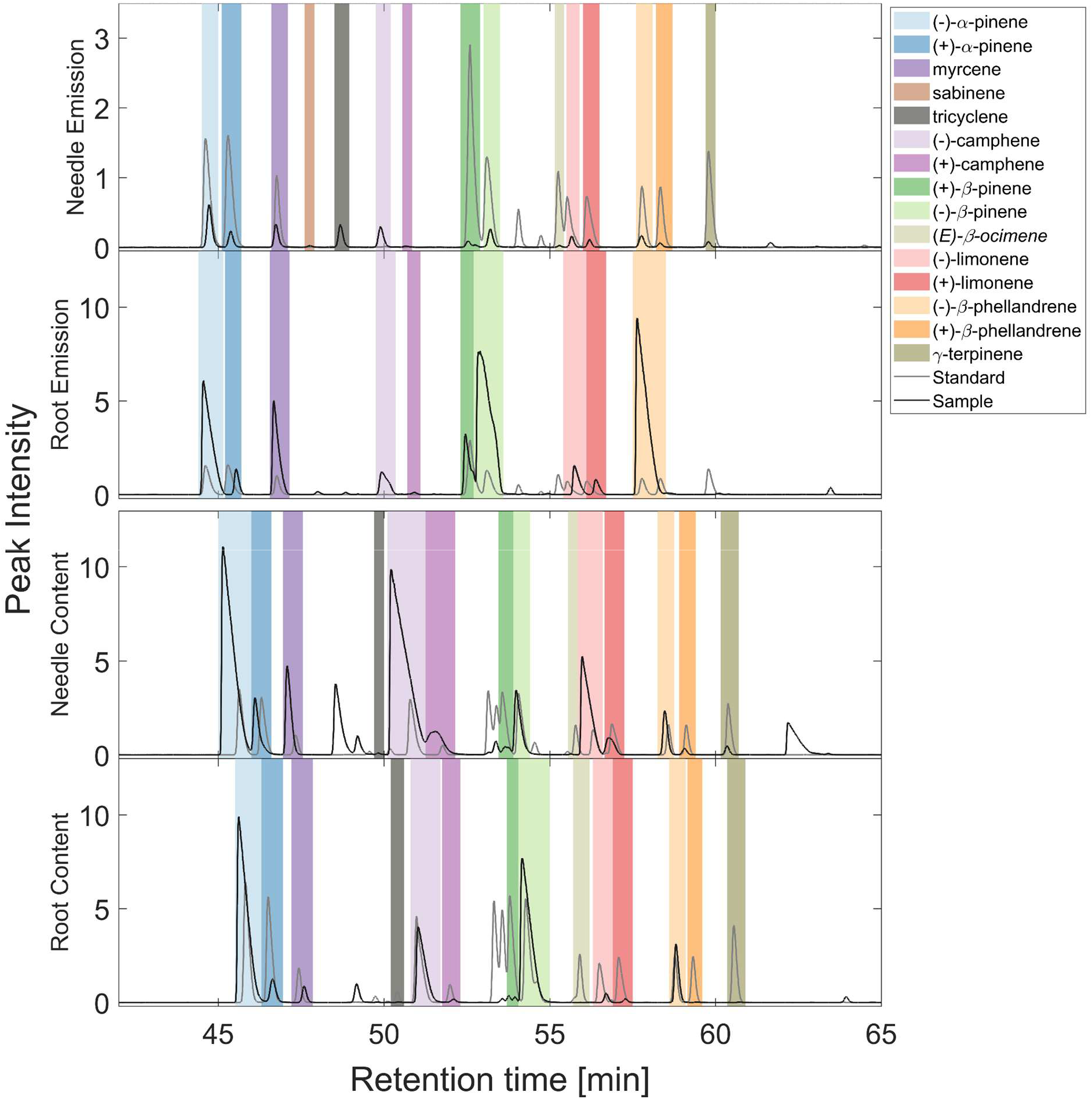
Gas chromatograms of mass 93 from a GC-MS of Tenax cartridges for emissions and methanol extracts on PDMS tubes of needle and root content of Norway spruce, and a standard mixture in hexane containing all chiral compounds. Peak intensity is displayed in 10^5^-steps. Colours correspond with the legend. Retention times partially vary between sample types due to adjustments in tuning and method. Identification was based on retention time and fragmentation pattern compared to mass-spectral NIST library (National Institute of Standards and Technology) and the standard. See **Fig. S2-16** for a detailed comparison of the GCMS spectra between the different sample types.

### Position-specific ^13^C2 pyruvate labelling

Isotopic tracer experiments are an effective tool to study the biosynthetic regulation of monoterpene storage and their emission rates. Pyruvate is a central intermediate and utilised as a precursor for various biosynthetic pathways in plant metabolism (Tcherkez *et al.*, 2005). For monoterpene biosynthesis, pyruvate can either enter the cytosolic mevalonate (MVA), or the plastidic methylerythritol 4-phosphate (MEP) pathway (Rohmer, 1999). In most reactions leading towards monoterpene biosynthesis, the C1 position of pyruvate is decarboxylated, whereas the C2 and C3 carbon atoms are integrated into the synthesized monoterpenes. Hence, to evaluate whether or not monoterpenes are synthesized *de novo* in roots and needles, we used ^13^C2-pyruvate in the labelling experiments. Experiments were performed on roots and branches of Norway spruce saplings after 21 days of the experiment on drought impacted and control plants to analyse the contribution of cytosolic intermediates to *de novo* synthesis of monoterpenes (**Fig. 1**). Branch labelling was performed based on previous studies (Fasbender *et al.*, 2018; Jardine *et al.*, 2014; Kreuzwieser *et al.*, 2021; Priault, Wegener and Werner, 2009; Tcherkez *et al.*, 2005; Tcherkez *et al.*, 2012)(**Fig. 1c**). One day before the experiments, branches were placed in the chambers to ensure equilibration and verify gas tightness. After 15 minutes of pre-measurements, the twig was cut close to the connected branch, placed under water and cut again to avoid embolism. 5 minutes after placement in water, the twig was placed in a tube containing 10 mM ^13^C2-labelled pyruvate (Cambridge Isotope Laboratories, Andover, MA, USA) solution of known weight for 80 minutes. At the end of the experiment, label uptake was determined by weighing the tubes again. Leaf areas were determined by GSA Image Analyzer v.4.1.8 (GSA GmbH, Rostock, Germany). Root labelling was developed based on the labelling procedure described by Honeker *et al.* (2022) and the online root bag sampling setup described by Lin, Owen and Peñuelas (2007) (**Fig. 1, 1d**). Roots were carefully excavated and placed into open-ended glass chambers (42 mm inner Ø, 125 mm height). The ends were sealed around the roots, using terostat (Terostat-II, Kahmann & Ellerbrock GmbH & Co. KG, Bielefeld, Germany) and PTFE foil. The tips of the roots were placed in water and emissions were monitored for 15 min before the water was exchanged with 80 mM ^13^C2-labelled pyruvate.

### Terpene sampling and GC-MS-C-IRMS analysis

VOCs from leaf and root chambers were collected at a flow rate of 120 ml min^-1^ for 60 minutes at the chamber outlet with glass thermodesorption tubes filled with Tenax TA before and at the end of pyruvate labelling. After the labelling experiment was conducted, labelled and unlabeled needles/roots were sampled, split into two parts and weighed. One part was dried (> 72h, 60°C) and weighed again to determine tissue water content. The other part was immediately flash frozen in liquid nitrogen and stored at -80°C. 40 mg frozen needle powder were transferred into 1 ml methanol and shaken at 1,400 rpm for 45 min at 30°C, followed by centrifugation for 5 min. To adsorb monoterpenes on polydimethylsiloxane (PDMS) tubes (for details, see Kallenbach *et al.*, 2014), 50 μl supernatant and five PDMS tubes were added to 950 μl demineralised H_2_O. For analysis on GC-C-IRMS, PDMS tubes were taken out after shaking (60 min, 30°C, 1400 rpm), dried with a lint-free tissue, and transferred to empty thermodesorption tubes (Gerstel, Germany). Due to lower monoterpene concentrations in root material, 80 mg were weighed into 1 ml of methanol, and 300 μl supernatant were added to 700 μl demineralized H_2_O instead.

For quantification and determination of ^13^C enrichment in emitted chiral monoterpenes, a GC-MS (7890B GC System & 5975C MSD, Agilent Technologie, Böblingen, Germany) coupled to a combustion interfaced isotope ratio mass spectrometer (C-IRMS, GC5 & Isoprime PrecisION, Elementar Analysesysteme GmbH, Langenselbold, Germany) was used. The thermodesorption tubes, either containing PDMS tubes for extracted samples, or Tenax for air samples, were heated to 220°C for 5 min in the TDU (thermodesorption unit, Gerstel, Germany) to desorb sampled terpenes, which were subsequently cryotrapped at -70°C in the CIS (cold injection system, Gerstel, Germany). Next, the CIS was heated to 240°C for 3 min to release the terpenes onto the GC separation column (beta-dex 120 Chirality, 60m x 250μm x 0.25μm, Supelco, USA) with helium. The separation column was initially heated to 45°C for 1 min, followed by an increase by 2°C min^-1^ to 60°C, then by 1°C min^-1^ to 150°C and finally by 3.5°C min^-1^ to 210°C. The gas stream was split behind the GC column, directing 10% of the gas stream towards the MSD, which was run in SIM mode to detect *m/z* 68, 93, 119 and 136 to identify sampled terpenes. 90% of the gas stream were directed towards the IRMS to measure ^13^C/^12^C ratios after passing the combustion interface to oxidize the terpenes to CO_2_ at 850°C, and a Nafion water trap to remove H_2_O. Spectral analysis was performed with the MassHunter software (Agilent Technologies, Böblingen, Germany). Compound identification was based on retention-times and fragmentation patterns found in the mass-spectral NIST library (National Institute of Standards and Technology), followed by manual control and reintegration of each individual peak. Identification was cross-compared to authentic standards that were also used for calibration ((−)-α-pinene, (+)-α-pinene, (−)-β-pinene, (+)-β-pinene, (−)-limonene, (+)-limonene, myrcene, (E)-β-ocimene, sabinene, camphene, β-phellandrene and α-phellandrene). For quantification of tricyclene and γ-terpinene, the calibration of (+)-β-pinene was used.

### Flux calculations

Molar flow *u* [mol s^-1^] was calculated as followed:

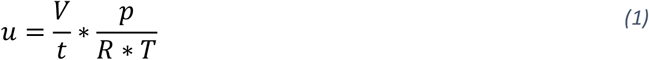

By taking the factor between gas volume *V* [m^3^] and time *t* [s], normalised to pressure *p* [Pa] and temperature *T* [K], and the universal gas constant *R* (8.31446 [J mol^-1^ K^-1^]). Assimilation rate *A* [μmol m^-2^ s^-1^] and transpiration rate *A* [mmol m^-2^ s^-1^] were determined after Caemmerer & Farquhar (1981). Water use efficiency (WUE), was calculated as the factor between assimilation and transpiration. BVOC fluxes *f*_*l*_ in [nmol m^-2^ s^-1^] were calculated from the VMR of the according BVOC from the empty chamber entrance of the leaf chamber *v*_*e*_ [ppb], and the VMR of the BVOC at the outlet of the leaf chamber *v*_*o*_ [ppb] using:

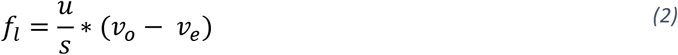

Isotopic composition of root CO_2_ *δ*^13^*C*_*r*_ [‰] (VPDB) was calculated using isotopic mass balance:

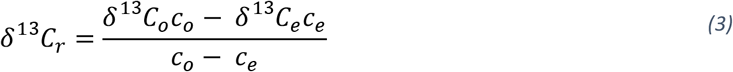

Where *δ*^13^*C*_*o*_ and *δ*^13^*C*_*e*_ are the delta values of the air at the chamber outlet, and entrance, respectively. The CO_2_ concentrations at the chamber outlet and entrance are described as *c*_*o*_ and *c*_*e*_.

### Statistical analysis

Matlab software version R2021b (MATLAB, 2021) was used for data processing, visualisation and statistical analysis. To determine significant differences between plant physiological parameters, VOC fluxes and monoterpene concentrations and composition in response to drought, the non-parametric Kruskal-Wallis test was used, followed by a post-hoc multiple comparison Tukey’s HSD test.

## Results

### Ecophysiological response to drought

Corresponding to the decline in soil VWC under drought to ∼28% from control conditions, assimilation, transpiration, δ^13^C of needle and root CO_2_ emissions, stomatal conductance, as well as root water content, were significantly affected (**Fig.3**). Assimilation and transpiration were halved in response to drought, thereby maintaining water use efficiency (WUE), while root respiration was not affected. On-line discrimination of ^13^CO_2_ by photosynthesis reduced by δ^13^C 0.5‰ and root CO_2_ emissions increased by δ^13^C 5‰ during drought. Root water content was reduced by one third compared to control conditions (**Fig.3**).

**Fig. 3.**
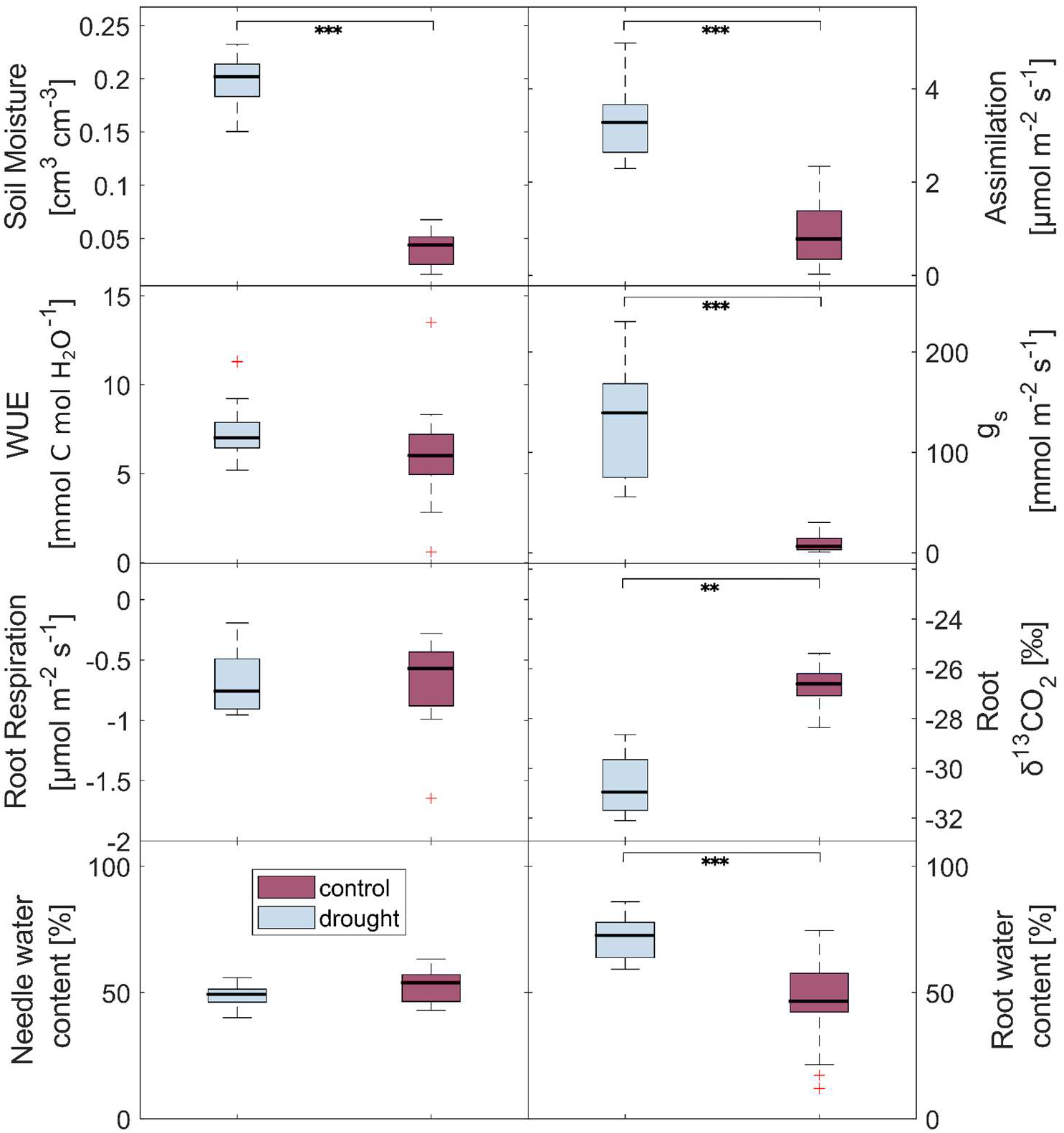
Ecophysiological response of *Picea abies* exposed to drought. Boxes indicate medians with 75^th^ and 25^th^ percentile as upper and lower limit, respectively (Control: needles n = 15, roots n = 10; Drought: needles n = 16, roots n = 15). WUE, water use efficiency; g_s_, stomatal conductance.

### Opposing drought responses of monoterpene content and emissions in both, needles and roots

Total monoterpene emissions of needles where significantly higher than emissions of roots in control (3-fold) and drought (2-fold) conditions, respectively (**Fig. 4a, S1**). Monoterpene content in needles was ten times higher than root monoterpene content under control, and seven times higher under drought conditions. Regarding the ratio between monoterpene emissions and tissue content, a significantly higher portion of monoterpenes was emitted per h by roots in comparison to needles, during control conditions (roots: 0.35‰; needles: 0.1‰), but not in drought conditions (roots: 0.05‰; needles: 0.045‰) (**Fig. 4b**). While monoterpene emissions from needles and roots were significantly reduced by drought to one third of control levels, monoterpene content was elevated by a factor of 1.5 in needles and 2.3 in roots, respectively (**Fig. 4a**).

**Fig. 4.**
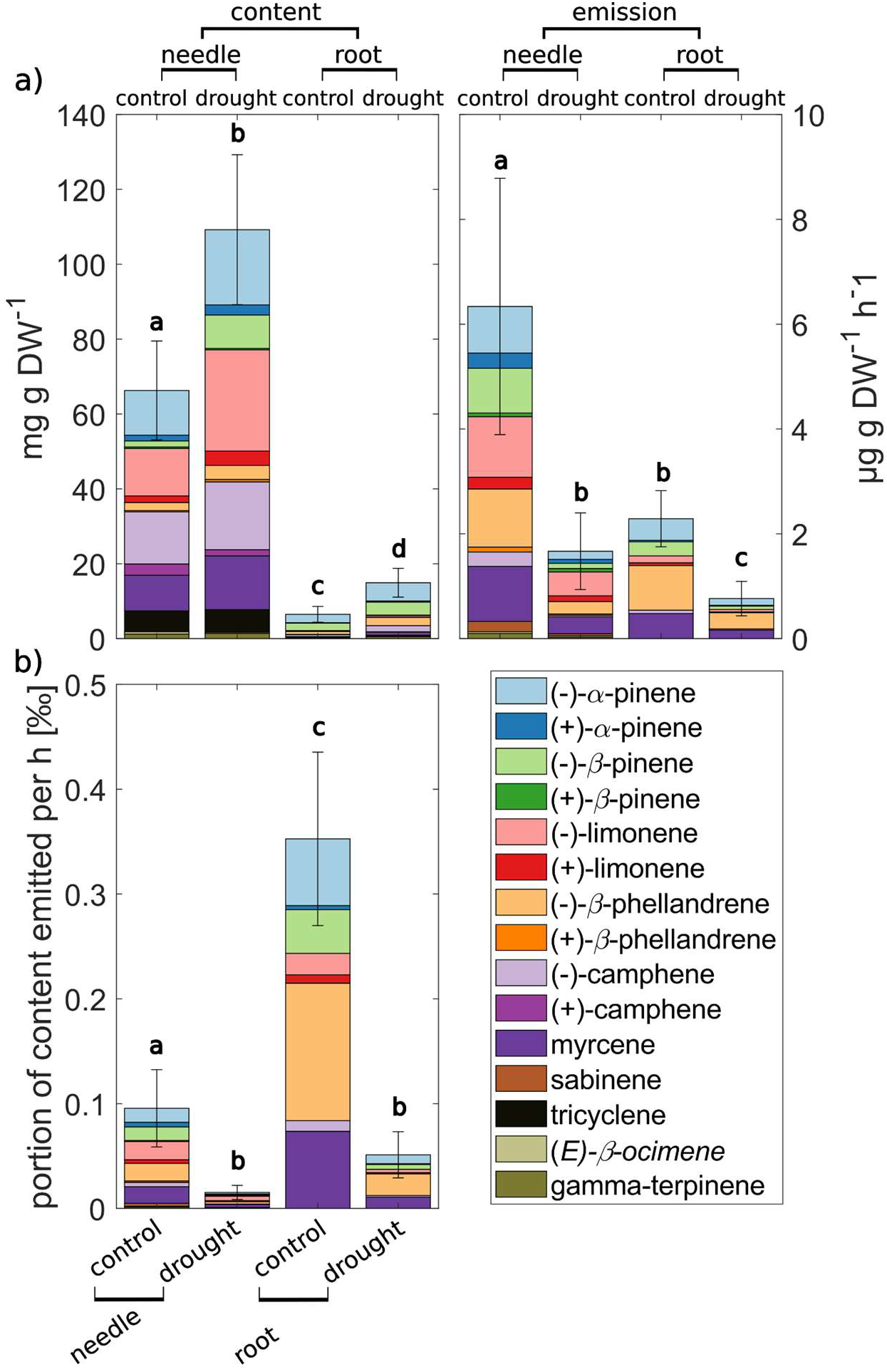
Speciation and quantification of chiral monoterpene emissions and content of needles and roots of *Picea abies* under control and drought conditions. a) Drought effect on the concentration of the 15 monoterpenes identified by GC-MS. Colours correspond with the legend in b). Error bars indicate standard deviation of total monoterpene concentration, significant differences (p < 0.05) are indicated by letters. Groups with different letters differ significantly in concentration. b) Portion of monoterpene content emitted per hour in %.

### Drought affects chiral monoterpene composition in needles but not in roots

Enantiomeric ratios were generally dominated by the (−)-enantiomers in both tissues (**Fig. 5, S1**). About 65% of total needle monoterpene emission and content were driven by (−)-enantiomers during control conditions. In roots the proportion of (−)-enantiomers was even higher with 75% and 85% in emissions and storage pools, respectively. In comparison, less than 5% of total monoterpene emissions and storage pools originated from (+)-enantiomers in roots and 10-15% in needle storage pools and emissions (**Fig. 5**). Interestingly, the ratio between (−)- and (+)-β-pinene significantly declined through drought in needle emissions, but significantly increased in needle content (**Fig 5b**). Ratios for other stereoisomers were not affected by drought.

**Fig. 5.**
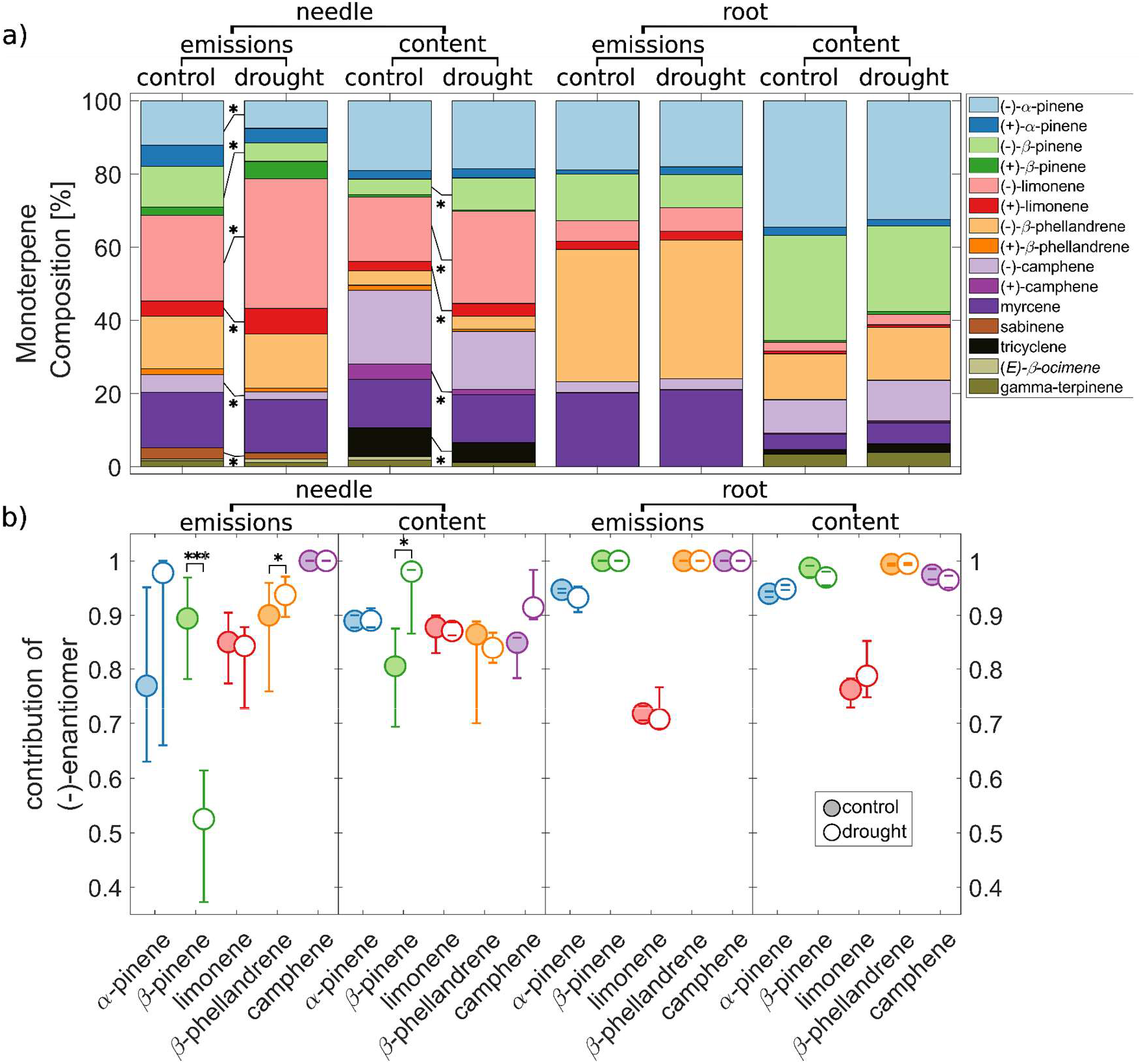
Composition of enantiomeric monoterpene emissions and content of needles and roots of *Picea abies* in response to drought. a) Drought effect on the composition of the 15 monoterpenes identified by GC-MS. Asterisks indicate significant differences in composition between control and drought phase (* p < 0.05). b) Contribution of the (−)-enantiomer to the overall concentrations of enantiomer couples in needle and root emissions and storage pools. Circles indicate medians with 75th and 25th percentile as upper and lower limit, respectively (Control: needle emissions n = 18, needle content n = 10, root emissions n = 11, root content n = 12; Drought: needle emissions n = 18, needle content n = 10, root emissions n = 17, root content n = 11). A value of 1 indicates that only the (−)-enantiomer is abundant. The colours correspond with the legend in a). Asterisks indicate significant differences between control and drought (* p < 0.05, *** p < 0.001).

Monoterpene composition differed between needles and roots (**Fig. 5**). Root monoterpene emissions and content were driven by (−)-α-pinene, (−)-β-pinene, (−)-β-phellandrene, (−)-camphene and myrcene, with minor contributions of (−)-limonene, and composition was not affected by drought. In contrast, (−)-limonene was the dominant monoterpene in needle emissions and content, and composition significantly changed in response to drought (**Fig. 5**). While the contribution of both enantiomers of limonene significantly increased in response to drought in needle emissions and content, contribution of the (−)-enantiomers of α-and β-pinene, as well as camphene to total needle monoterpene emissions significantly declined. In contrast, the contribution of (−)-α-pinene to total needle monoterpene content was not affected by drought.

### δ^13^C2-labelled pyruvate is not incorporated into chiral monoterpene pools of needles and roots and root emissions

We investigated the incorporation of ^13^C into chiral monoterpenes to evaluate the contribution of freshly assimilated carbon to chiral monoterpene content and their emissions, via GC-MS-C-IRMS, by labelling the branches and roots with ^13^C2-pyruvate during control and drought conditions. Pyruvate is a central metabolite in plant metabolism and is an intermediate for the synthesis of monoterpenes and several other VOCs. Under control conditions, significant amounts of ^13^C of ^13^C2-pyruvate were emitted as CO_2_, and were incorporated into acetaldehyde and ethanol emitted from roots and needles and, in the case of needle emissions, also acetone (**Fig. S1**). We observed significant amounts of ^13^C2-label being incorporated into the needle emissions of (−)-limonene, (+)-limonene and (+)-alpha-pinene under control conditions. However, we could not detect any label incorporation into needle and root pools nor into root emissions (**Fig. 6**). In response to drought, label incorporation in monoterpenes almost completely declined in needle emissions and no label was detected in any other treatment (data not shown).

**Fig. 6.**
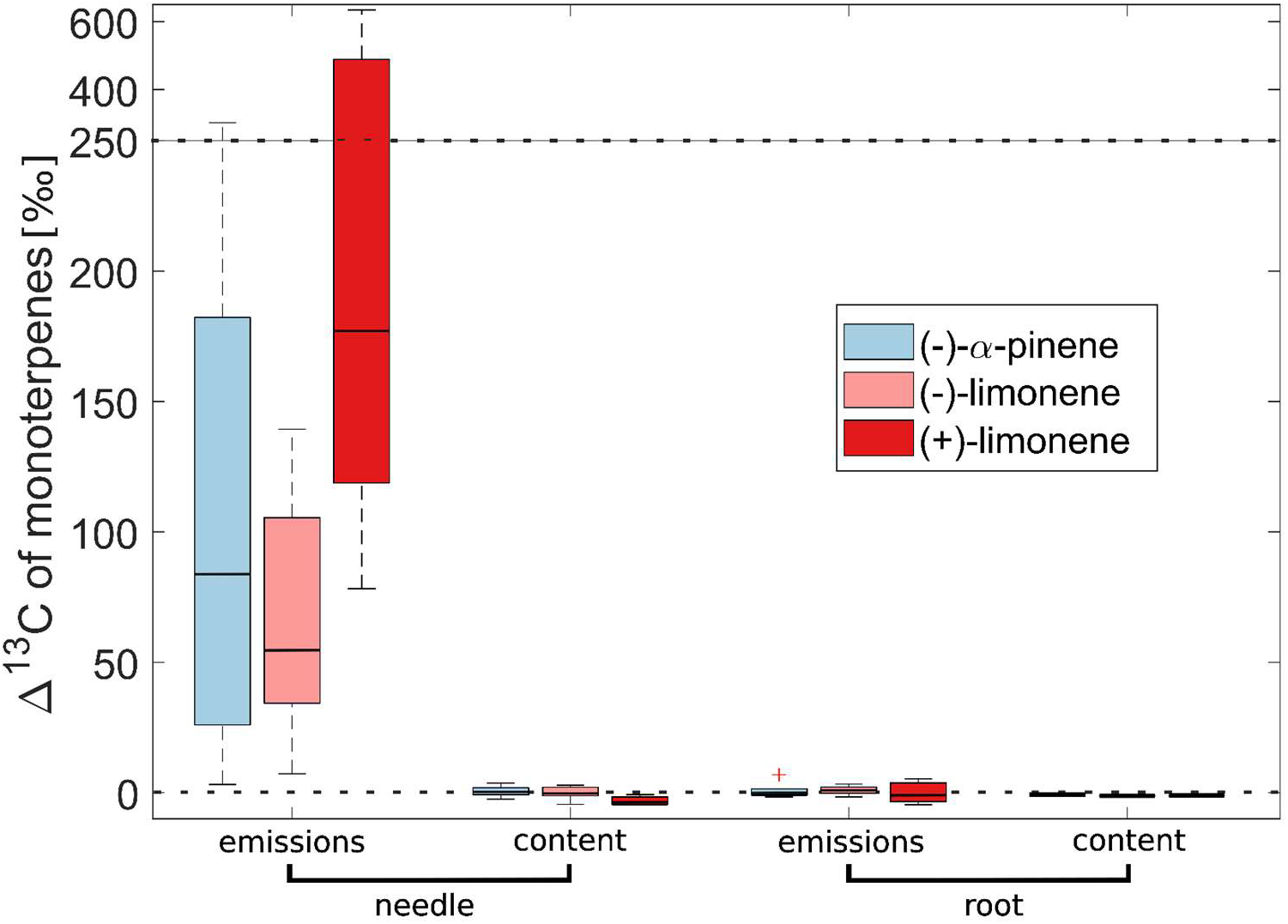
Label incorporation into chiral monoterpenes emitted and stored by *Picea abies* after ^13^C2-pyruvate labelling during control conditions, presented as change Δ in δ^13^C of monoterpenes before the labelling compared to after labelling. Boxes indicate medians with 75th and 25th percentile as upper and lower limit, respectively (needle emissions n = 7, needle content n = 6, root emissions n = 7, root content n = 6). Values above 0 indicate ^13^C-incorporation after labelling. The y-axis is split for better visualisation.

## Discussion

Chiral monoterpenes emitted by plants above- and belowground shape the chemical landscape of many ecosystems, significantly affecting the interactions of plants with their environment (Billings and Cameron, 1984; de los Santos and Wolf, 2020; Dudareva *et al.*, 2013; Schulz-Bohm *et al.*, 2018). Interactions, for example with herbivores, strongly depend on signals indicating plant physiological status which are affected by environmental stressors, such as drought (Dobbertin *et al.*, 2007; Holopainen *et al.*, 2018; Lehmanski *et al.*, 2023). Our results demonstrate how drought unequally affects biosynthesis, emissions and pool size of above- and belowground chiral monoterpenes, outlining the complexity of effects on plant metabolism by drought. The observed decline in chiral monoterpene emissions of both, needles and roots of Norway spruce, while simultaneously increasing storage pools (**Fig. 7**), indicates that emissions from storage pools are not only constrained by temperature, which will be further discussed below.

**Fig. 7.**
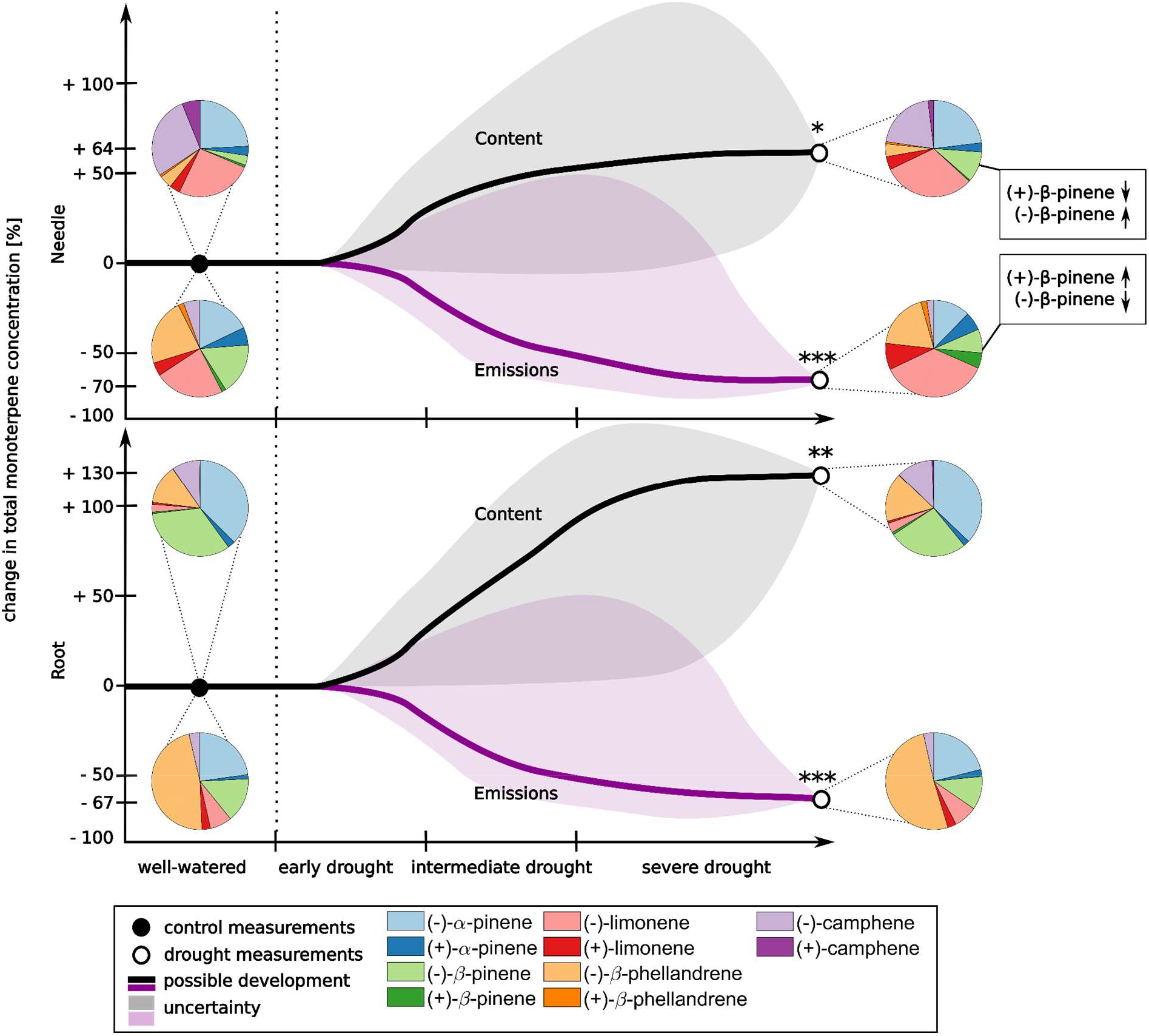
Schematic description of relative changes in needle and root monoterpene content and emissions in response to drought. Lines describe a possible development of concentrations throughout the drought. Shaded areas represent uncertainties based on previously published findings. Asterisks indicate significant differences between control and drought measurements of cumulative monoterpene concentrations (* p < 0.05, ** p < 0.01, *** p < 0.001). Pie charts show chiral monoterpene composition during control and drought conditions for each treatment.

### Tissue-specific chiral monoterpene composition indicates tissue-specific chiral monoterpene functions and biosynthetic controls

Chiral monoterpene composition differed between needles and roots, and between emissions and content (**Fig. 5, 7**). Chiral monoterpene composition of needles changed in response to drought: contribution of both enantiomers of limonene significantly increased in needle emissions and storage pools, whereas contribution of other (−)-enantiomers declined in emissions, but increased or remained stable in storage pools. In contrast, chiral monoterpene composition remained stable in roots in response to drought. The most notable difference between needle and root monoterpene composition was the low contribution of both limonene enantiomers and high contribution of (−)-enantiomers of pinenes in roots. Interestingly, (−)-enantiomers dominated in both, needles and roots, whereas in Scots pine, branch emissions were dominated by (+)-enantiomers in another study (Staudt *et al.*, 2019).

The distinct variation in chiral monoterpene composition between needles and roots suggests a linkage between monoterpene allocation and functioning, and it is likely that different mixtures of monoterpenes vary in their attractive/deterrent properties for different organisms (Lehmanski *et al.*, 2023). Thus, tissue-specific chiral monoterpene composition and quantity might have developed as a consequence of tissue-specific pest-species (Borg-Karlson *et al.*, 1993).

Stereoisomers of limonene possess insecticidal properties that deter bark beetles (Fang *et al.*, 2021; Zhao *et al.*, 2010), highlighting the relevance of high aboveground emissions of limonene, even under drought (**Fig. 4**). The observed shift in the chiral ratio of β-pinene in emissions and content of needles likely affected biological properties of the monoterpene mixture. The opposing trends between emissions and content, with the portion of (+)-β-pinene increasing in emissions, but decreasing in content, could indicate active control of (+)-β-pinene emissions from needle content in response to drought. On the one hand the observed shift in chiral composition could increase deterring potential (Da Rivas Silva *et al.*, 2012). On the other hand, these shifts could also be distinct cues, for e.g. primary attraction of bark beetles (Lehmanski *et al.*, 2023), which should be further investigated via isolated attraction studies with different non-racemic chiral monoterpene mixtures. However, by accounting for the contribution of compounds with demonstrated antimicrobial/insecticidal properties (both limonene enantiomers, (+)-α-pinene and (+)-β-pinene), the portion of antimicrobial/insecticidal monoterpenes in needle emissions increased in response to drought from 34% to 50% and in needle storage pools from 26% to 32%, whereas the antimicrobial/insecticidal portion remained unchanged in roots (∼7-10%) (**Fig. 5**). Therefore, antimicrobial/insecticidal properties might be of higher relevance for aboveground emissions compared to belowground emission, especially during drought stress. With the bark beetle *Ips typographus* being the most destructive pest species of Norway spruce, it seems plausible that aboveground emissions are optimised for defensive properties. In addition, the OH reactivity of aboveground emissions increased in response to drought (Daber *et al. in review*). Thus, compositional changes might indicate an optimisation for protection from oxidative and herbivore stress at low emission (Jardine *et al.*, 2017). This could also explain why compositional changes only occurred aboveground. To better understand host selection by bark beetle individuals, future studies should aim to investigate whether drought also affects the composition and magnitude of emissions and storage pools of chiral monoterpenes of stems, the site of attack.

Our novel approach to investigate belowground emissions of chiral monoterpene composition could provide important cues for our understanding of the interactions between plant roots and other organisms. Chiral monoterpene composition likely affects chemotaxis, i.e. bacterial migration towards roots (Schulz-Bohm *et al.*, 2018). Soilborne bacteria and fungi are known to be able to utilise various monoterpenes as carbon and energy source (Harder and Probian, 1995; Marmulla and Harder, 2014; Seubert, 1960), which is affected by their toxicity and hence, chiral composition (Da Rivas Silva *et al.*, 2012; van Vuuren and Viljoen, 2007). However, chemotaxis by and utilisation of chiral monoterpenes affects the interaction with both, beneficial (Keel, 1992;Kleinheinz *et al.*, 1999) and pathogenic (Tan, Day and Cadwallader, 1998) microbes, resulting in a trade-off for plants between attraction of symbionts and deterrence of pathogens (el Zahar Haichar *et al.*, 2014). Based on their strong antimicrobial and antimycotic effects (van Vuuren and Viljoen, 2007), we would have expected high belowground emissions of both limonene enantiomers. Conversely, limonene emissions were comparatively low in roots. However, roots exude a plethora of other, non-volatile chemicals into the rhizosphere, including antimicrobial compounds (Baetz and Martinoia, 2014). Hence, belowground olfactory cues of chiral monoterpenes might be rather released as signalling molecules for mutualistic interactions (Ali, Alborn and Stelinski, 2010; Rasmann *et al.*, 2005).

Notably, tissue-specific variations in monoterpene storage composition have been reported in the past for Norway spruce and were speculated to occur due to tissue-specific differences in biosynthetic pathways (Borg-Karlson *et al.*, 1993; Persson *et al.*, 1996; Persson, Borg-Karlson and Norin, 1993). At least four multi-product- and one single-product mTPS produce the diverse mixture of chiral monoterpenes emitted and stored by Norway spruce (Fäldt *et al.*, 2003; Martin, Fäldt and Bohlmann, 2004). Among these mTPS, limonene synthase (PaTPS-Lim) mainly produces the enantiomers of limonene, while the dominant products of pinene synthase (PaTPS-Pin) are the (−)-enantiomers of β-pinene, α -pinene and β-phellandrene (Martin, Fäldt and Bohlmann, 2004). Based on these studies, our results indicate that mTPS activity may be tissue-specific, with limonene synthase being only active in aboveground tissue. However, the low biosynthetic rates for (+)-enantiomers reported by these two studies, neither match with our observed chiral ratios in needle tissue nor with those from previous studies (Borg-Karlson *et al.*, 1993; Persson, Borg-Karlson and Norin, 1993). It is therefore likely that other, currently unidentified mTPS contribute to the complex monoterpene mixtures produced by Norway spruce. Alternatively, or additionally, product rates of these mTPS might be affected by different external factors, e.g. the physiologic status of the cell. In this case, the shifts of chiral ratios in response to drought, observed for needle content and emissions (**Fig. 5**) could also be explained. However, further studies, combining measurements of chiral monoterpene emissions and storage content with analyses of mTPS expression levels under different environmental stressors will be necessary to unravel the underlying biosynthetic controls driving these processes. RNA analyses were not included in the funding for the presented work but could be usefully incorporated in future studies

The discrepancies observed between the monoterpene composition of emissions and storage content in both, needles and roots (**Fig. 5**) could be due to differences in mTPS composition and activity in parenchymatic cells, compared to mTPS composition within leucoplasts, fuelling storage pool refill (Cheniclet and Carde, 1985). Though, it remains open why monoterpene composition of emissions does not match the composition of storage compartments in needles and roots under drought, when *de novo* synthesis declines (**Fig. 5, 6**), which will be further discussed in the following.

### Monoterpene emissions are controlled independently from storage pools

We observed a corresponding decline between photosynthesis, transpiration and *de novo* biosynthesis and emission of monoterpenes in response to drought (**Fig. 3, 4a, 5, 7, S1**). In contrast, monoterpene pool size increased during drought. Thus, the portion of monoterpenes emitted relative to the total monoterpene content in both, needles and roots significantly declined under drought (**Fig. 4b**). About 30% of total needle monoterpene emissions from Norway spruce are synthesised *de novo* in undisturbed conditions (Ghirardo *et al.*, 2010). Therefore, the observed decline in needle emissions by more than 60% under drought can only partially be explained by the decline in *de novo* synthesis.

Hence, monoterpene emissions from storage pools also declined in response to drought, even though total monoterpene content increased. The non-specific storage of monoterpenes in mesophyll cells is too small to sustain emissions throughout long-term stress (Niinemets and Reichstein, 2002). Thus, the remaining emissions under drought are maintained by release from resin ducts and associated cells. It is generally assumed that monoterpene emissions from storage pools are released passively into the intercellular air space in dependence to temperature, even after *de novo* synthesis diminishes (Fischbach *et al.*, 2002; Gershenzon, McConkey and Croteau, 2000). With increasing storage pool concentrations in response to drought and identical temperatures compared to control conditions, monoterpene emissions from resin ducts should therefore increase, or at least remain of similar magnitude compared to the control phase, but instead they decline (**Fig. 4**). Notably, the observed decline in stomatal conductance (**Fig. 3**) during drought cannot explain the lower emission rates, since, in general, the emission of monoterpenes is not affected by stomatal conductance, as long as foliage intercellular air space monoterpene concentrations are maintained, e.g. by a constant release from storage compartments (Loreto *et al.*, 1996; Niinemets, Loreto and Reichstein, 2004). Consequently, monoterpene emissions decline in response to drought due to diminishing monoterpene concentrations within the intercellular air space. Based on our findings, we conclude that needle monoterpenes are not passively emitted from storage compartments, with composition and magnitude being determined by total monoterpene content of these structures, but that their release is rather subject to further controls. We assume that needle-internal monoterpene release from high-content resin ducts and associated cells to the needle surface is disrupted under drought. It should be noted that the discrepancies we observed between monoterpene composition of storage compartments and emissions (**Fig. 5**) cannot be explained by physical properties of different monoterpenes, such as boiling point differences; In this case emissions from compounds with relatively low boiling points (α-pinene, β-pinene) from storage compartments should have been higher, compared to compounds with relatively high boiling points (β-phellandrene), but this was not the case. Alternatively, active transport mechanisms could be involved, as proposed for monoterpene transport between biosynthetic cells and storage compartments in peppermint plants (McCaskill, Gershenzon and Croteau, 1992). More recently, Chang *et al.* (2023) identified two ABC-transporters in the orchid *Phaleonopsis bellina* regulating floral emissions of monoterpenes. These findings encourage considerations of a transporter-based regulation of monoterpene emissions in response to environmental stressors. However, we can only speculate whether this disruption is actively controlled or a consequence of drought stress, and our findings give rise for further investigations of needle internal regulation of monoterpene release in response to drought.

Little is known about root monoterpene emissions and even less about their response to drought (Lin, Owen and Peñuelas, 2007). Similar to the observed dynamics in needles, we also observed opposing trends between monoterpene content and emission in roots in response to drought, as well as compositional differences (**Fig. 5, Fig. 7**). Our results of the isotopic tracer experiments indicate that no significant amounts of *de novo* biosynthesis of monoterpenes take place inside the roots. Even though the isotopic tracer was taken up by the roots and utilised for biosynthesis and emission of other VOCs, monoterpenes were not labelled (**Fig. 6, S2, S3**). Nonetheless, it is likely that monoterpenes can be produced *de novo* in roots, as high concentrations of root TPSs were found in other studies (Zhou and Pichersky, 2020). Chen *et al.* (2004) showed that 1,8-cineole is directly produced and emitted by the roots of *Arabidopsis*, but not accumulated in root tissue. It is assumed that monoterpene biosynthesis and emission is driven by the root cortex or surrounding mucilage (Lin, Owen and Peñuelas, 2007). Therefore, selective monoterpene transport from resin ducts towards the root cortex, or biosynthesis of monoterpenes via transported isoprenoid intermediates from shoots (Burlat *et al.*, 2004) into the root cortex, could explain our findings. Nonetheless, emissions could also be affected by changes in exudate composition that have been observed in response to drought in various plant species (Williams and de Vries, 2020).

In summary, chiral monoterpene composition is tissue-specific and likely related to tissue-specific functioning. Monoterpene emissions do not correspond to total monoterpene content, even when *de novo* synthesis diminishes, but seem to be subject to further constraints. These results are important to consider in modelling approaches, as potential increases in monoterpene emissions in response to rising temperatures (Guenther *et al.*, 2012), could be buffered by control of storage release in response to drought. Our findings imply important ramifications for plant insect interactions and the effect of shifts in chiral compositions (as observed for β-pinene) should be further investigated, e.g. in primary attraction studies with bark beetles.

## Acknowledgements

This research was funded by the European Research Council (ERC consolidator grant 647008 to CW) and the German Research Foundation (ECOSENSE SFB 1537). We gratefully acknowledge help from Philipp Nolte and Chidubem Anene with data collection, Monika Eiblmeier for technical assistance in GC-MS/IRMS analyses, Michael Rienks for logistical help and Eva Schottmüller for assisting with gardening the plants.

## Author Contributions

LED, CW and JK planned and designed the research. LED and MM performed experiments and collected data. LED, CW and JK analysed and interpreted the data. LED wrote the manuscript with input from all co-authors.

## Open Research

The data that support the findings of this study are available from the corresponding author upon reasonable request.

## Notes

### Competing Interest Statement

The authors have declared no competing interest.

## Reference list

Aggarwal, K.K. et al. (2002) ‘Antimicrobial activity profiles of the two enantiomers of limonene and carvone isolated from the oils of Mentha spicata and Anethum sowa’, Flavour and Fragrance Journal, 17(1), pp. 59–63. doi: 10.1002/ffj.1040

Ali, J.G., Alborn, H.T. and Stelinski, L.L. (2010) ‘Subterranean herbivore-induced volatiles released by citrus roots upon feeding by Diaprepes abbreviatus recruit entomopathogenic nematodes’, Journal of Chemical Ecology, 36(4), pp. 361–368. doi: 10.1007/s10886-010-9773-7

Aubourg, S., Lecharny, A. and Bohlmann, J. (2002) ‘Genomic analysis of the terpenoid synthase (AtTPS) gene family of Arabidopsis thaliana’, Molecular Genetics and Genomics, 267(6), pp. 730– 745. doi: 10.1007/s00438-002-0709-y

Baetz, U. and Martinoia, E. (2014) ‘Root exudates: the hidden part of plant defense’, Trends in Plant Science, 19(2), pp. 90–98. doi: 10.1016/j.tplants.2013.11.006

Bernard-Dagan, C. (1988) ‘Seasonal Variations in Energy Sources and Biosynthesis of Terpenes in Maritime Pine’, in Mechanisms of Woody Plant Defenses Against Insects: Search for Pattern.: Springer New York, pp. 93–116. Available at: https://link.springer.com/chapter/10.1007/978-1-4612-3828-7_5.

Billings, R.F. and Cameron, R.S. (1984) ‘Kairomonal Responses of Coleoptera, Monochamus titillator (Cerambycidae), Thanasimus dubius (Cleridae), and Temnochila virescens (Trogositidae), to Behavioral Chemicals of Southern Pine Bark Beetles (Coleoptera: Scolytidae)’, Environmental Entomology, 13(6), pp. 1542–1548. doi: 10.1093/ee/13.6.1542

Borg-Karlson, A.K. et al. (1993) ‘Enantiomeric composition of monoterpene hydrocarbons in different tissues of Norway Spruce, Picea abies (L) Karst. A multi-dimensional gas chromatography study’, Acta Chemica Scandinavia, 47, pp. 138–144.

Bos, R. et al. (2002) ‘Volatile components from Anthriscus sylvestris (L.) Hoffm.’, Journal of Chromatography a, 966(1-2), pp. 233–238. doi: 10.1016/S0021-9673(02)00704-5

Burlat, V. et al. (2004) ‘Co-expression of three MEP pathway genes and geraniol 10-hydroxylase in internal phloem parenchyma of Catharanthus roseus implicates multicellular translocation of intermediates during the biosynthesis of monoterpene indole alkaloids and isoprenoid-derived primary metabolites’, The Plant Journal, 38(1), pp. 131–141. doi: 10.1111/j.1365-313X.2004.02030.x

Byron, J. et al. (2022) ‘Chiral monoterpenes reveal forest emission mechanisms and drought responses’, Nature, 609(7926), pp. 307–312. doi: 10.1038/s41586-022-05020-5

Caemmerer, S. von and Farquhar, G.D. (1981) ‘Some relationships between the biochemistry of photosynthesis and the gas exchange of leaves’, Planta, 153(4), pp. 376–387. doi: 10.1007/BF00384257

Chang, Y.-L. et al. (2023) ‘PbABCG1 and PbABCG2 transporters are required for the emission of floral monoterpenes in Phalaenopsis bellina’, The Plant Journal : for Cell and Molecular Biology, 114(2), pp. 279–292. doi: 10.1111/tpj.16133

Chen, F. et al. (2004) ‘Characterization of a root-specific Arabidopsis terpene synthase responsible for the formation of the volatile monoterpene 1,8-cineole’, Plant Physiology, 135(4), pp. 1956–1966. doi: 10.1104/pp.104.044388

Chen, F. et al. (2011) ‘The family of terpene synthases in plants: a mid-size family of genes for specialized metabolism that is highly diversified throughout the kingdom’, The Plant Journal, 66(1), pp. 212–229. doi: 10.1111/j.1365-313X.2011.04520.x

Cheng, Z. et al. (2017) ‘The research of genetic toxicity of β-phellandrene’, Environmental Toxicology and Pharmacology, 54, pp. 28–33. doi: 10.1016/j.etap.2017.06.011

Cheniclet, C. and Carde, J.P. (1985) ‘Presence of leucoplasts in secretory cells and of monoterpenes in the essential oil: a correlative study’, Israel Journal of Plant Sciences, 34(2-4), pp. 219–238. doi: 10.1080/0021213X.1985.10677023

Da Rivas Silva, A.C. et al. (2012) ‘Biological activities of α-pinene and β-pinene enantiomers’, Molecules, 17(6), pp. 6305–6316. doi: 10.3390/molecules17066305

de los Santos, Z.A. and Wolf, C. (2020) ‘Optical Terpene and Terpenoid Sensing: Chiral Recognition, Determination of Enantiomeric Composition and Total Concentration Analysis with Late Transition Metal Complexes’, Journal of the American Chemical Society, 142(9), pp. 4121–4125. doi: 10.1021/jacs.9b13910

Delory, B.M. et al. (2016) ‘Root-emitted volatile organic compounds: can they mediate belowground plant-plant interactions?’ Plant and Soil, 402(1-2), pp. 1–26.

Dobbertin, M. et al. (2007) ‘Linking increasing drought stress to Scots pine mortality and bark beetle infestations’, The Scientific World Journal, 7, pp. 231–239.

Dudareva, N. et al. (2013) ‘Biosynthesis, function and metabolic engineering of plant volatile organic compounds’, New Phytologist, 198(1), pp. 16–32.

el Zahar Haichar, F. et al. (2014) ‘Root exudates mediated interactions belowground’, Soil Biology and Biochemistry, 77, pp. 69–80. doi: 10.1016/j.soilbio.2014.06.017

Fäldt, J. et al. (2003) ‘Traumatic resin defense in Norway spruce (Picea abies): Methyl jasmonate-induced terpene synthase gene expression, and cDNA cloning and functional characterization of (+)-3-carene synthase’, Plant molecular biology, 51(1), pp. 119–133.

Fang, J.X. et al. (2021) ‘Functional investigation of monoterpenes for improved understanding of the relationship between hosts and bark beetles’, Journal of Applied Entomology, 145(4), pp. 303–311. doi: 10.1111/jen.12850

Fasbender, L. et al. (2018) ‘Real-time carbon allocation into biogenic volatile organic compounds (BVOCs) and respiratory carbon dioxide (CO2) traced by PTR-TOF-MS, 13CO2 laser spectroscopy and 13C-pyruvate labelling’, PloS One, 13(9), e0204398.

Fischbach, R.J. et al. (2002) ‘Seasonal pattern of monoterpene synthase activities in leaves of the evergreen tree Quercus ilex’, Physiologia Plantarum, 114(3), pp. 354–360.

Flores, H.E., Vivanco, J.M. and Loyola-Vargas, V.M. (1999) ‘‘Radicle’biochemistry: the biology of root-specific metabolism’, Trends in Plant Science, 4(6), pp. 220–226.

Gershenzon, J., McConkey, M.E. and Croteau, R.B. (2000) ‘Regulation of monoterpene accumulation in leaves of peppermint’, Plant Physiology, 122(1), pp. 205–214. doi: 10.1104/pp.122.1.205

Ghirardo, A. et al. (2010) ‘Determination of de novo and pool emissions of terpenes from four common boreal/alpine trees by 13CO2 labelling and PTR-MS analysis’, Plant, Cell & Environment, 33(5), pp. 781–792.

Guenther, A.B. et al. (1993) ‘Isoprene and monoterpene emission rate variability: Model evaluations and sensitivity analyses’, Journal of Geophysical Research, 98(D7), p. 12609. doi: 10.1029/93JD00527

Guenther, A.B. et al. (2012) ‘The Model of Emissions of Gases and Aerosols from Nature version 2.1 (MEGAN2. 1): an extended and updated framework for modeling biogenic emissions’, Geoscientific Model Development, 5(6), pp. 1471–1492.

Hachlafi, N.E.L. et al. (2023) ‘In Vitro and in Vivo Biological Investigations of Camphene and Its Mechanism Insights: A Review’, Food Reviews International, 39(4), pp. 1799–1826. doi: 10.1080/87559129.2021.1936007

Hans, J. et al. (2004) ‘Cloning, characterization, and immunolocalization of a mycorrhiza-inducible 1-deoxy-d-xylulose 5-phosphate reductoisomerase in arbuscule-containing cells of maize’, Plant Physiology, 134(2), pp. 614–624. doi: 10.1104/pp.103.032342

Harder, J. and Probian, C. (1995) ‘Microbial degradation of monoterpenes in the absence of molecular oxygen’, Applied and Environmental Microbiology, 61(11), pp. 3804–3808. doi: 10.1128/aem.61.11.3804-3808.1995

Holopainen, J.K. et al. (2018) ‘Climate Change Effects on Secondary Compounds of Forest Trees in the Northern Hemisphere’, Frontiers in Plant Science, 9, p. 1445. doi: 10.3389/fpls.2018.01445

Honeker, L.K. et al. (2022) ‘Elucidating Drought-Tolerance Mechanisms in Plant Roots through 1H NMR Metabolomics in Parallel with MALDI-MS, and NanoSIMS Imaging Techniques’, Environmental Science & Technology, 56(3), pp. 2021–2032. doi: 10.1021/acs.est.1c06772

Janson, R.W. (1993) ‘Monoterpene emissions from Scots pine and Norwegian spruce’, Journal of Geophysical Research: Atmospheres, 98(D2), pp. 2839–2850.

Jardine, K.J. et al. (2014) ‘Phytogenic biosynthesis and emission of methyl acetate’, Plant, Cell and Environment, 37(2), pp. 414–424. doi: 10.1111/pce.12164

Jardine, K.J. et al. (2017) ‘Monoterpene ‘thermometer’ of tropical forest-atmosphere response to climate warming’, Plant, Cell and Environment, 40(3), pp. 441–452. doi: 10.1111/pce.12879

Kallenbach, M. et al. (2014) ‘A robust, simple, high-throughput technique for time-resolved plant volatile analysis in field experiments’, The Plant Journal : for Cell and Molecular Biology, 78(6), pp. 1060–1072. doi: 10.1111/tpj.12523

Keel, C. (1992) ‘Suppression of Root Diseases by Pseudomonas fluorescens CHA0: Importance of the Bacterial Secondary Metabolite 2,4-Diacetylphloroglucinol’, Molecular Plant-Microbe Interactions, 5(1), p. 4. doi: 10.1094/MPMI-5-004

Keeling, C.I. et al. (2011) ‘Transcriptome mining, functional characterization, and phylogeny of a large terpene synthase gene family in spruce (Picea spp.)’, BMC Plant Biology, 11(1), p. 43. doi: 10.1186/1471-2229-11-43

Kleinheinz, G.T. et al. (1999) ‘Characterization of alpha-pinene-degrading microorganisms and application to a bench-scale biofiltration system for VOC degradation’, Archives of Environmental Contamination and Toxicology, 37(2), pp. 151–157. doi: 10.1007/s002449900500

Kreuzwieser, J. et al. (2021) ‘Drought affects carbon partitioning into volatile organic compound biosynthesis in Scots pine needles’, New Phytologist, 232(5), pp. 1930–1943.

Krupa, S. and Fries, N. (1971) ‘Studies on ectomycorrhizae of pine. I. Production of volatile organic compounds’, Canadian Journal of Botany, 49(8), pp. 1425–1431. doi: 10.1139/b71-200

Lehmanski, L.M.A. et al. (2023) ‘Addressing a century-old hypothesis - do pioneer beetles of Ips typographus use volatile cues to find suitable host trees?’ New Phytologist, 238(5), pp. 1762–1770. doi: 10.1111/nph.18865

Lewinsohn, E. et al. (1993) ‘Oleoresinosis in Grand Fir (Abies grandis) saplings and mature trees (modulation of this wound response by light and water stresses)’, Plant Physiology, 101(3), pp. 1021–1028.

Lin, C., Owen, S.M. and Peñuelas, J. (2007) ‘Volatile organic compounds in the roots and rhizosphere of Pinus spp’, Soil Biology and Biochemistry, 39(4), pp. 951–960.

Loreto, F. et al. (1996) ‘Influence of Environmental Factors and Air Composition on the Emission of alpha-Pinene from Quercus ilex Leaves’, Plant Physiology, 110(1), pp. 267–275. doi: 10.1104/pp.110.1.267

Loreto, F. et al. (1998) ‘Measurement of isoprenoid content in leaves of Mediterranean Quercus spp. by a novel and sensitive method and estimation of the isoprenoid partition between liquid and gas phase inside the leaves’, Plant Science, 136(1), pp. 25–30.

Loreto, F. et al. (2000) ‘Emission and content of monoterpenes in intact and wounded needles of the Mediterranean Pine, Pinus pinea’, Functional Ecology, 14(5), pp. 589–595. doi: 10.1046/j.1365-2435.2000.t01-1-00457.x

Marmulla, R. and Harder, J. (2014) ‘Microbial monoterpene transformations-a review’, Frontiers in Microbiology, 5, p. 346. doi: 10.3389/fmicb.2014.00346

Martin, D.M., Fäldt, J. and Bohlmann, J. (2004) ‘Functional characterization of nine Norway spruce TPS genes and evolution of gymnosperm terpene synthases of the TPS-d subfamily’, Plant Physiology, 135(4), pp. 1908–1927.

MATLAB (2021) version 9.11 (R2021b). Natick, Massachusetts: The MathWorks Inc.

McCaskill, D., Gershenzon, J. and Croteau, R. (1992) ‘Morphology and monoterpene biosynthetic capabilities of secretory cell clusters isolated from glandular trichomes of peppermint (Mentha piperita L.)’, Planta, 187(4), pp. 445–454. doi: 10.1007/BF00199962

Neirotti, E., Moscatelli, M. and Tiscornia, S. (1996) ‘Antimicrobial activity of the limonene’, Arquivos de Biologia e Tecnologia, 39, pp. 233–238.

Niinemets, Ü., Loreto, F. and Reichstein, M. (2004) ‘Physiological and physicochemical controls on foliar volatile organic compound emissions’, Trends in Plant Science, 9(4), pp. 180–186.

Niinemets, Ü. and Reichstein, M. (2002) ‘A model analysis of the effects of nonspecific monoterpenoid storage in leaf tissues on emission kinetics and composition in Mediterranean sclerophyllous Quercus species’, Global Biogeochemical Cycles, 16(4), 57–1.

Norin, T. (1996) ‘Chiral chemodiversity and its role for biological activity. Some observations from studies on insect/insect and insect/plant relationships’, Pure and Applied Chemistry, 68(11), pp. 2043–2049. doi: 10.1351/pac199668112043

Persson, M. et al. (1996) ‘Relative amounts and enantiomeric compositions of monoterpene hydrocarbons in xylem and needles of Picea abies’, Phytochemistry, 42(5), pp. 1289–1297. doi: 10.1016/0031-9422(96)00119-7

Persson, M., Borg-Karlson, A.K. and Norin, T. (1993) ‘Enantiomeric composition of six chiral monoterpene hydrocarbons in different tissues of Picea abies’, Phytochemistry, 33(2), pp. 303– 307.

Phillips, M.A., Savage, T.J. and Croteau, R. (1999) ‘Monoterpene synthases of loblolly pine (Pinus taeda) produce pinene isomers and enantiomers’, Archives of Biochemistry and Biophysics, 372(1), pp. 197–204. doi: 10.1006/abbi.1999.1467

Priault, P., Wegener, F. and Werner, C. (2009) ‘Pronounced differences in diurnal variation of carbon isotope composition of leaf respired CO2 among functional groups’, New Phytologist, 181(2), pp. 400–412. doi: 10.1111/j.1469-8137.2008.02665.x

Rasmann, S. et al. (2005) ‘Recruitment of entomopathogenic nematodes by insect-damaged maize roots’, Nature, 434(7034), pp. 732–737. doi: 10.1038/nature03451

Rohmer, M. (1999) ‘The discovery of a mevalonate-independent pathway for isoprenoid biosynthesis in bacteria, algae and higher plants’, Natural Product Reports, 16(5), pp. 565–574. doi: 10.1039/a709175c

Rufino, A.T. et al. (2014) ‘Anti-inflammatory and chondroprotective activity of (+)-α-pinene: structural and enantiomeric selectivity’, Journal of Natural Products, 77(2), pp. 264–269. doi: 10.1021/np400828x

Schulz-Bohm, K. et al. (2018) ‘Calling from distance: attraction of soil bacteria by plant root volatiles’, The ISME Journal, 12(5), pp. 1252–1262. doi: 10.1038/s41396-017-0035-3

Schürmann, W. et al. (1993) ‘Emission of biosynthesized monoterpenes from needles of Norway Spruce’, Naturwissenschaften, 80(6), pp. 276–278. doi: 10.1007/BF01135913

Seubert, W. (1960) ‘Degradation of isoprenoid compounds by micro-organisms. I. Isolation and characterization of an isoprenoid-degrading bacterium, Pseudomonas citronellolis n. sp’, Journal of Bacteriology, 79(3), pp. 426–434. doi: 10.1128/jb.79.3.426-434.1960

Shao, M. et al. (2001) ‘Volatile organic compound emissions from Scots pine: mechanisms and description by algorithms’, Journal of Geophysical Research: Atmospheres, 106(D17), pp. 20483– 20491.

Sjödin, K. et al. (1996) ‘Enantiomeric compositions of monoterpene hydrocarbons in different tissues of four individuals of Pinus sylvestris’, Phytochemistry, 41(2), pp. 439–445.

Staudt, M. et al. (1997) ‘Seasonal and diurnal patterns of monoterpene emissions from Pinus pinea (L.) under field conditions’, Atmospheric Environment, 31, pp. 145–156. doi: 10.1016/s1352-2310(97)00081-2

Staudt, M. et al. (2019) ‘Compartment specific chiral pinene emissions identified in a Maritime pine forest’, The Science of the Total Environment, 654, pp. 1158–1166. doi: 10.1016/j.scitotenv.2018.11.146

Steeghs, M. et al. (2004) ‘Proton-transfer-reaction mass spectrometry as a new tool for real time analysis of root-secreted volatile organic compounds in Arabidopsis’, Plant Physiology, 135(1), pp. 47–58. doi: 10.1104/pp.104.038703

Stotzky, G. and Schenck, S. (1976) ‘Volatile organic compounds and microorganisms’, CRC Critical Reviews in Microbiology, 4(4), pp. 333–382. doi: 10.3109/10408417609102303

Tan, Q., Day, D.F. and Cadwallader, K.R. (1998) ‘Bioconversion of (R)-(+)-limonene by P. digitatum (NRRL 1202)’, Process Biochemistry, 33(1), pp. 29–37. doi: 10.1016/S0032-9592(97)00048-4

Tcherkez, G. et al. (2005) ‘In vivo respiratory metabolism of illuminated leaves’, Plant Physiology, 138(3), pp. 1596–1606. doi: 10.1104/pp.105.062141

Tcherkez, G. et al. (2012) ‘Respiratory carbon fluxes in leaves’, Current Opinion in Plant Biology, 15(3), pp. 308–314. doi: 10.1016/j.pbi.2011.12.003

Trapp, S. and Croteau, R.B. (2001) ‘Defensive resin biosynthesis in conifers’, Annual Review of Plant Biology, 52(1), pp. 689–724.

Unsicker, S.B., Kunert, G. and Gershenzon, J. (2009) ‘Protective perfumes: the role of vegetative volatiles in plant defense against herbivores’, Current Opinion in Plant Biology, 12(4), pp. 479–485. doi: 10.1016/j.pbi.2009.04.001

van Vuuren, S.F. and Viljoen, A.M. (2007) ‘Antimicrobial activity of limonene enantiomers and 1,8-cineole alone and in combination’, Flavour and Fragrance Journal, 22(6), pp. 540–544. doi: 10.1002/ffj.1843

Wichtmann, E.M. and Stahl-Biskup, E. (1987) ‘Composition of the essential oils from caraway herb and root’, Flavour and Fragrance Journal, 2(2), pp. 83–89. doi: 10.1002/ffj.2730020207

Williams, A. and de Vries, F.T. (2020) ‘Plant root exudation under drought: implications for ecosystem functioning’, New Phytologist, 225(5), pp. 1899–1905. doi: 10.1111/nph.16223

Williams, J. et al. (2011) ‘The summertime Boreal forest field measurement intensive (HUMPPA-COPEC-2010): an overview of meteorological and chemical influences’, Atmospheric Chemistry and Physics, 11(20), pp. 10599–10618. doi: 10.5194/acp-11-10599-2011

Yassaa, N. and Williams, J. (2007) ‘Enantiomeric monoterpene emissions from natural and damaged Scots pine in a boreal coniferous forest measured using solid-phase microextraction and gas chromatography/mass spectrometry’, Journal of Chromatography a, 1141(1), pp. 138–144. doi: 10.1016/j.chroma.2006.12.006

Zhao, T. et al. (2010) ‘The influence of Ceratocystis polonica inoculation and methyl jasmonate application on terpene chemistry of Norway spruce, Picea abies’, Phytochemistry, 71(11-12), pp. 1332–1341. doi: 10.1016/j.phytochem.2010.05.017

Zhou, F. and Pichersky, E. (2020) ‘The complete functional characterisation of the terpene synthase family in tomato’, New Phytologist, 226(5), pp. 1341–1360.

