## Supplementary Information for "Chiral monoterpene dynamics of shoots and roots of Norway spruce in response to drought"

L. Erik Daber[1, *], Jürgen Kreuzwieser[1], Mirjam Meischner[1], Jonathan Williams[2], Christiane Werner[1]

[1]Chair of Ecosystem Physiology, Albert-Ludwigs-Universität Freiburg, Georges-Köhler-Allee 053/054, 79110 Freiburg, Germany

[2]Max Planck Institute for Chemistry, Hahn-Meitner-Weg 1, 55128 Mainz, Germany

*Author for correspondence:

*Lars Erik Daber*

**

**Contents:**

**Materials and methods** for calibration of the PTR-TOF-MS used to measure fluxes of selected VOCs

**Figure S1** Label incorporation into $CO_2$ and selected VOCs by *Picea abies* via $^{13}$C2-pyruvate labelling

**Figure S2-S16** GCMS spectrum of identified chiral monoterpenes from different samples of *Picea abies* and a Standard.

**Materials and methods** for calibration of the PTR-TOF-MS used to measure fluxes of selected VOCs

Calibration was performed using a liquid calibration unit (LCU, Ionicon Analytic GmbH, Innsbruck, Austria) in dependence of humidity and stepwise dilution of a multi-component calibration mixture in nitrogen (Apel Riemer Environmental Inc., USA). Volume mixing ratios (VMR, [ppbv]) were determined via mass-dependent transmission calculation followed by mass scale calibration, using PTRwid software (version 003 08-11-2020, (Holzinger, 2015)) after the procedure described by Holzinger *et al.* (2019).

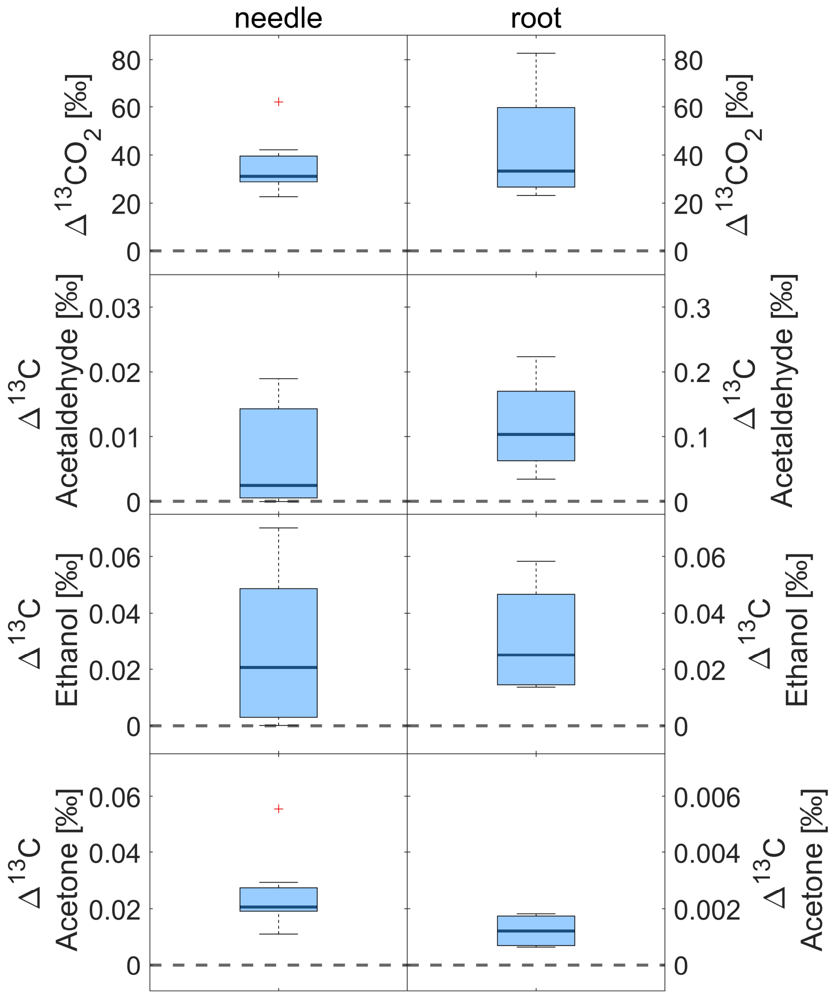


**Fig. S1** Label incorporation into $CO_2$ and selected VOCs by *Picea abies* after $^{13}$C2-pyruvate labelling, presented as difference Δ in $\delta^{13}C$ before and after the labelling. Boxes indicate medians with 75th and 25th percentile as upper and lower limit, respectively (needles n = 7, roots n = 7). Values above 0 indicate elevated $\delta^{13}C$ values after labelling. Label incorporation was measured via Delta Ray IRIS for $\delta^{13}C$ of $CO_2$ and by PTR-ToF-MS for VOCs.

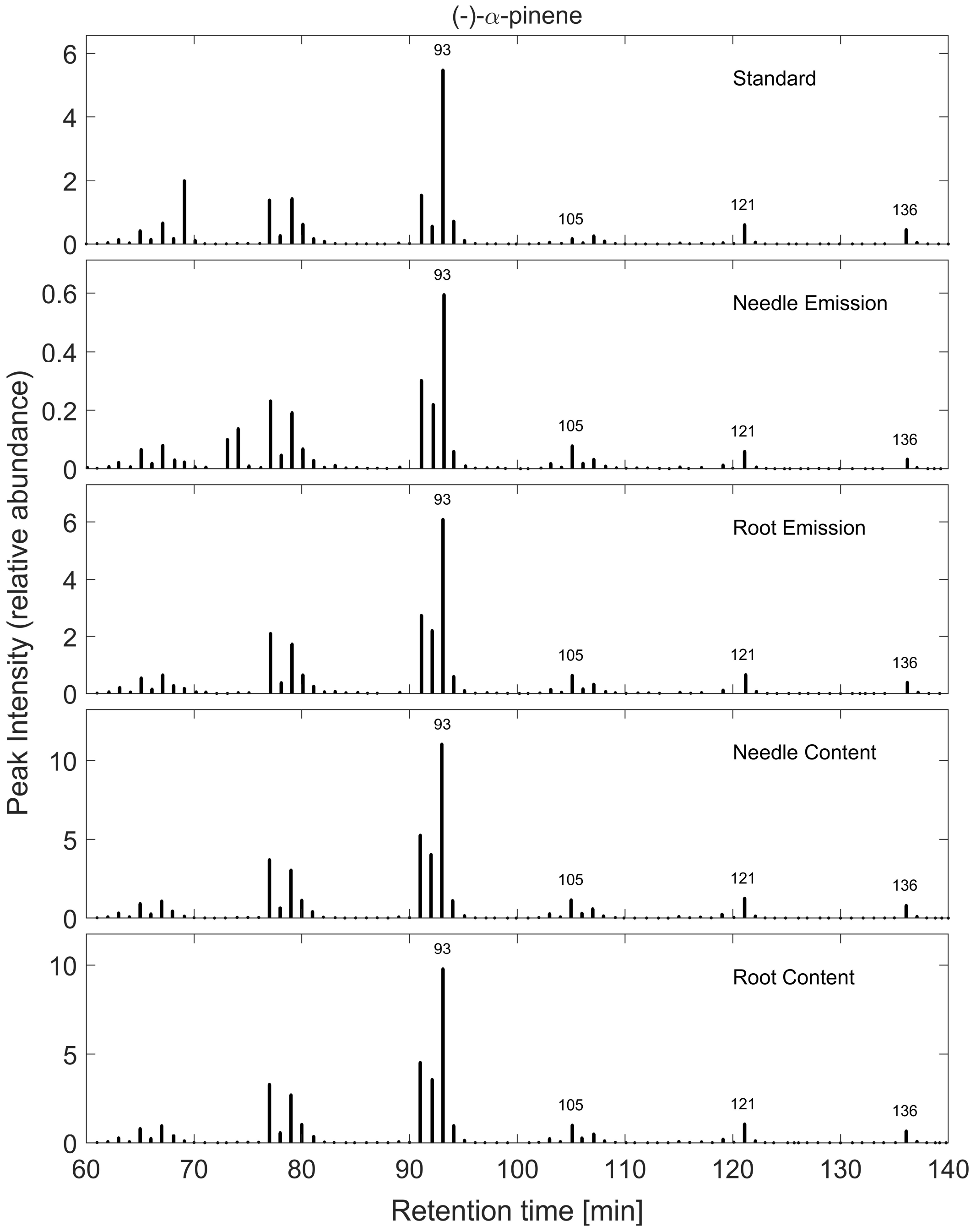


**Fig. S2** GCMS spectrum of (-)-α-pinene from different samples of *Picea abies* and a Standard. Peak intensity is displayed in $10^5$-steps.

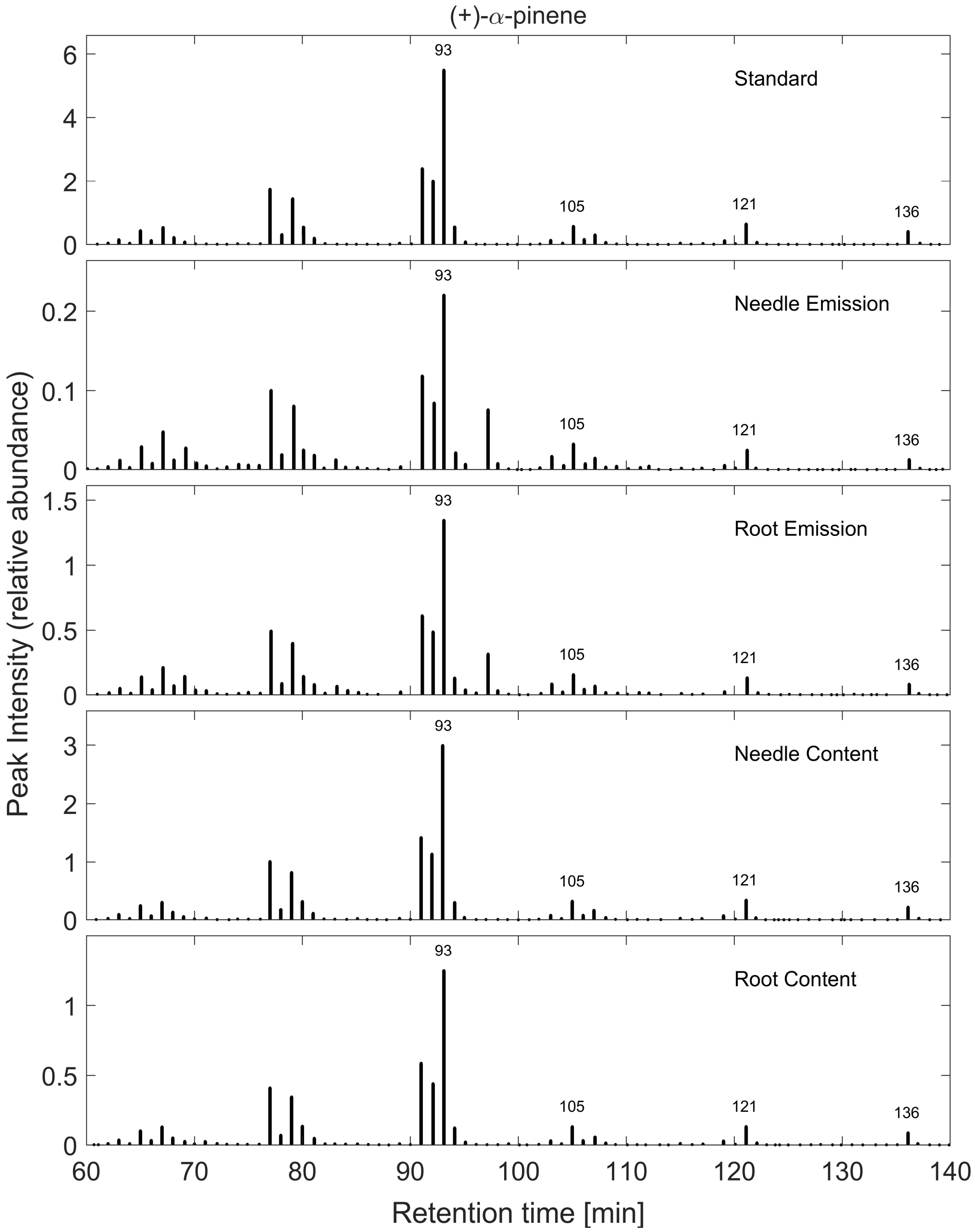


**Fig. S3** GCMS spectrum of (+)-α-pinene from different samples of *Picea abies* and a Standard. Peak intensity is displayed in $10^5$-steps.

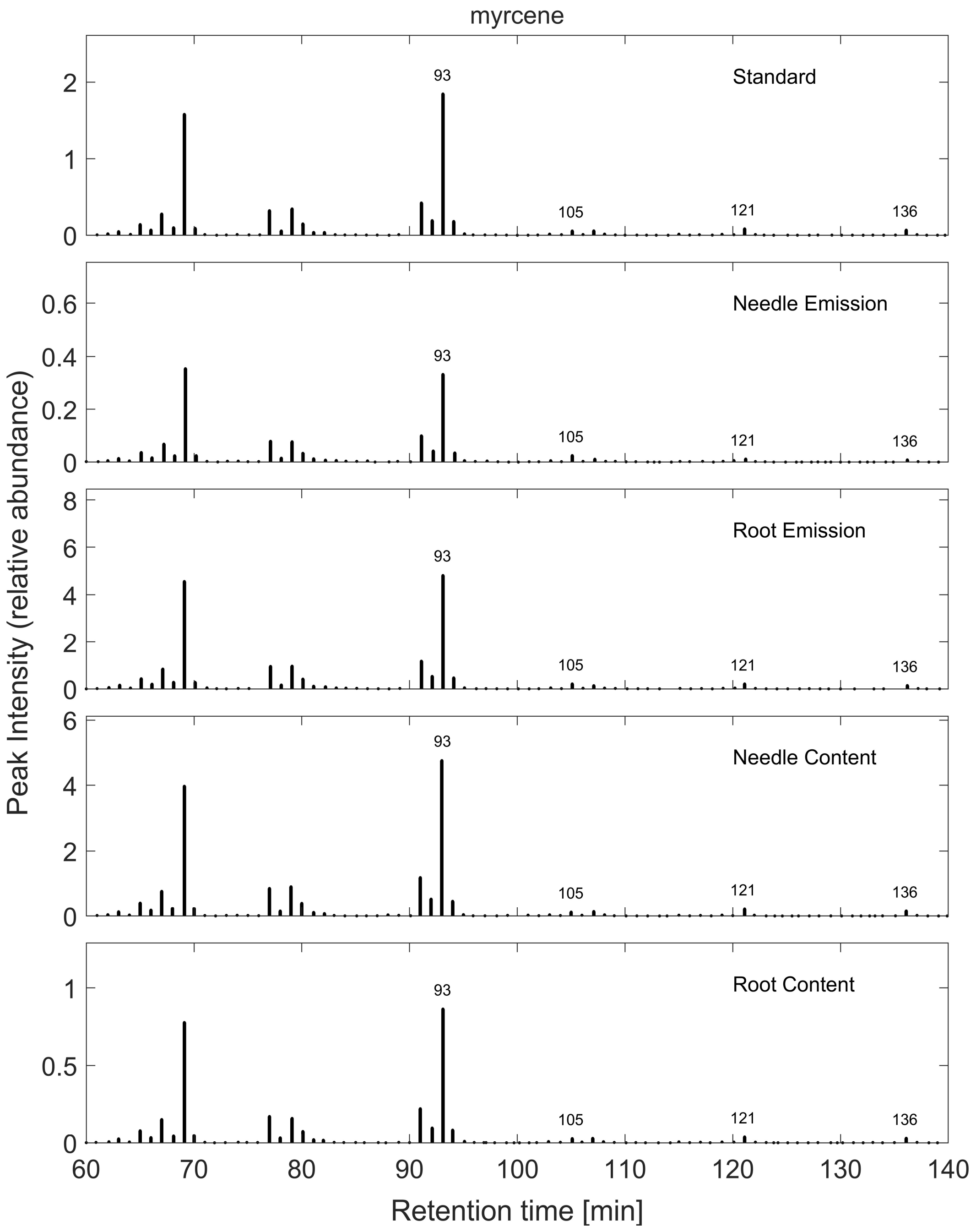


**Fig. S4** GCMS spectrum of myrcene from different samples of *Picea abies* and a Standard. Peak intensity is displayed in $10^5$-steps.

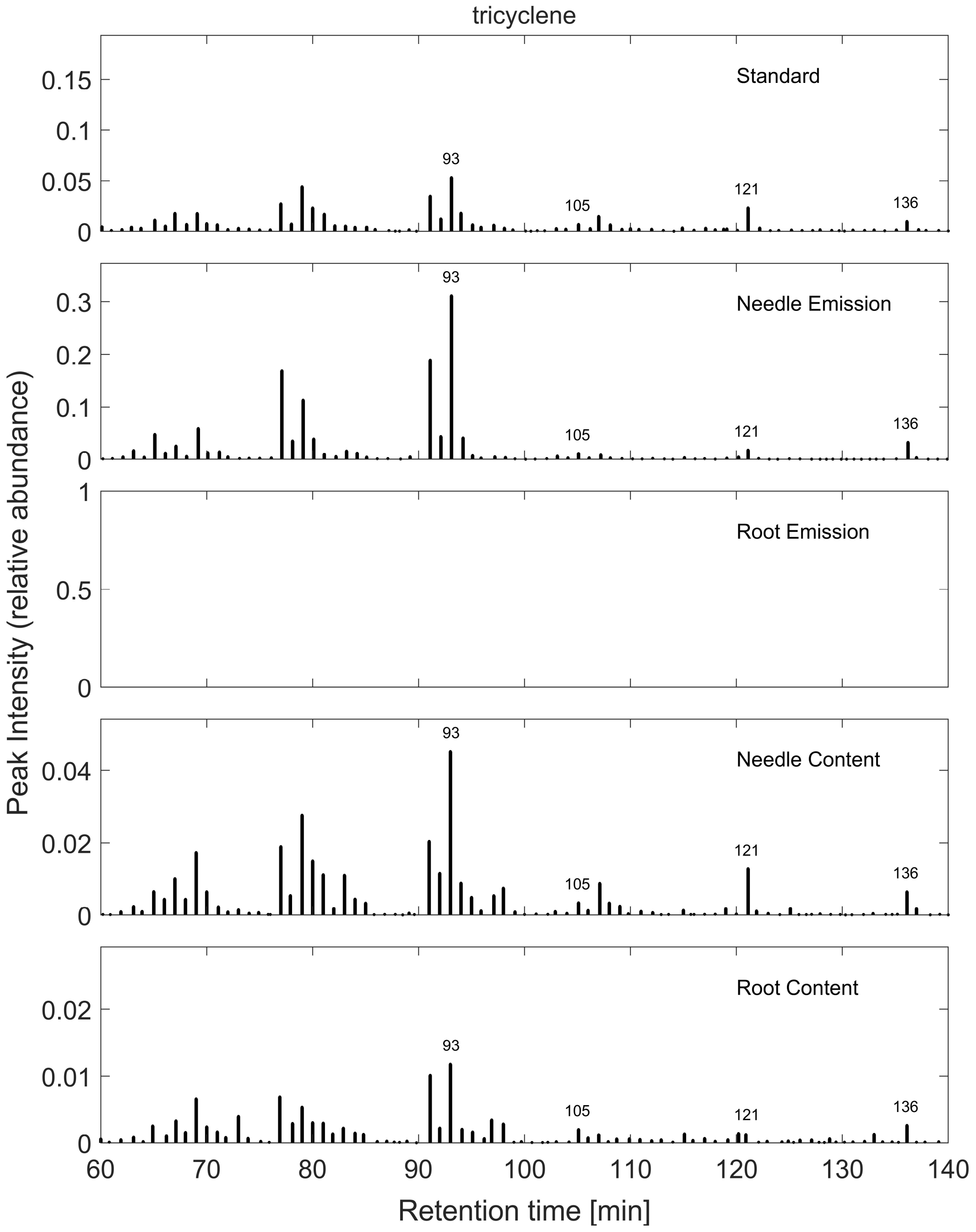


**Fig. S5** GCMS spectrum of tricyclene from different samples of *Picea abies* and a Standard. Peak intensity is displayed in $10^5$-steps. Empty graphs indicate that the terpene was not identified in the sample.

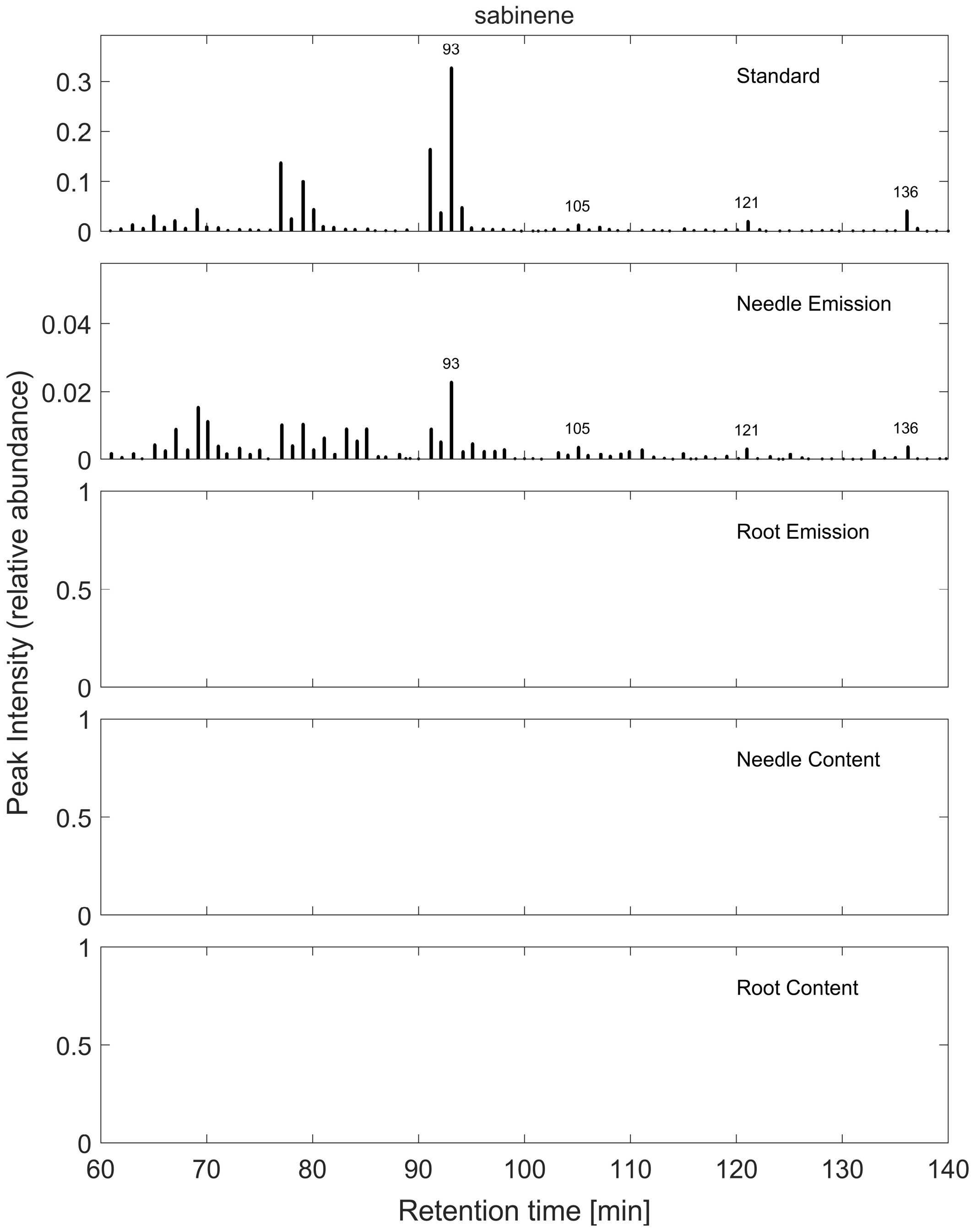


**Fig. S6** GCMS spectrum of sabinene from different samples of *Picea abies* and a Standard. Peak intensity is displayed in $10^5$-steps. Empty graphs indicate that the terpene was not identified in the sample.

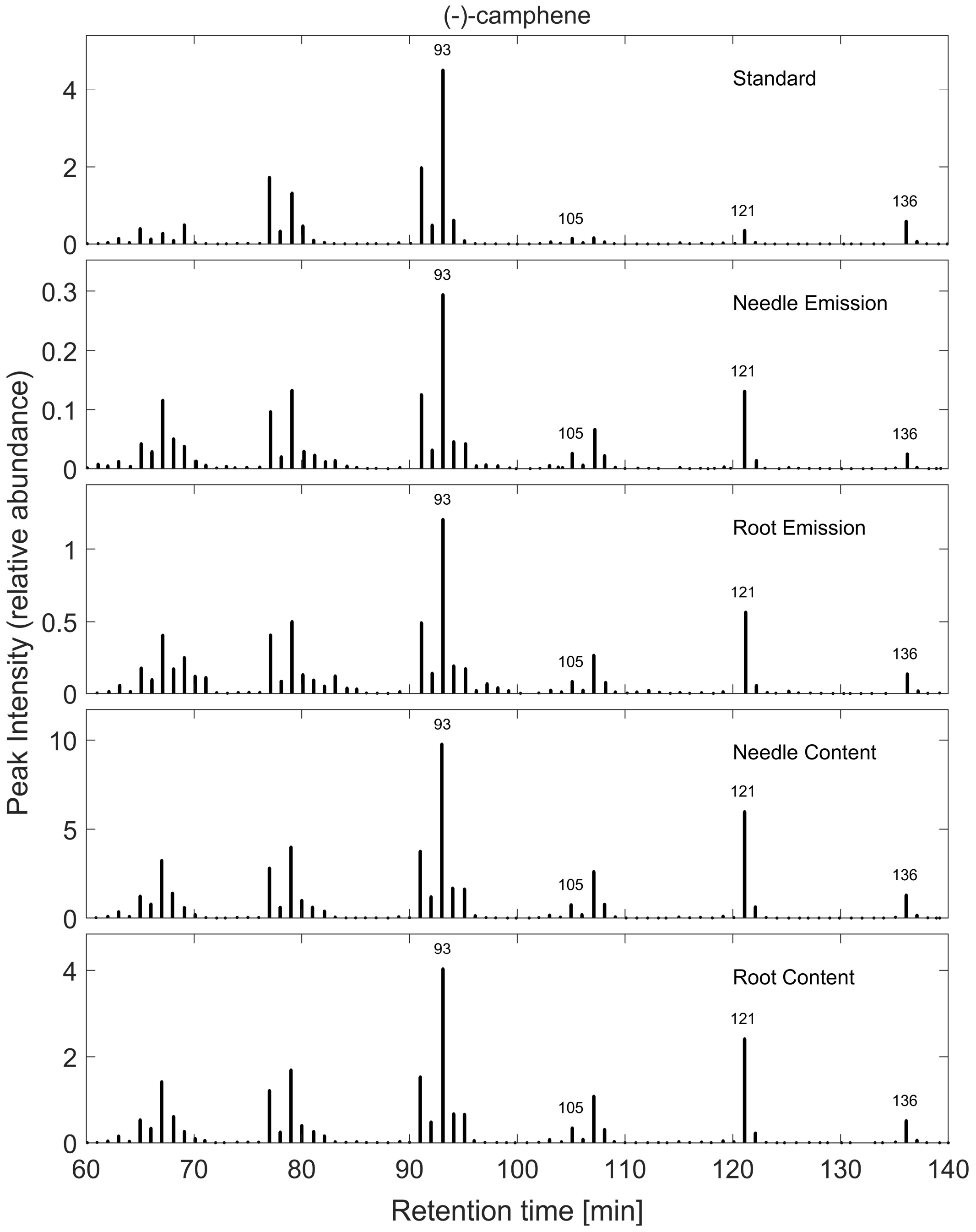


**Fig. S7** GCMS spectrum of (-)-camphene from different samples of *Picea abies* and a Standard. Peak intensity is displayed in $10^5$-steps.

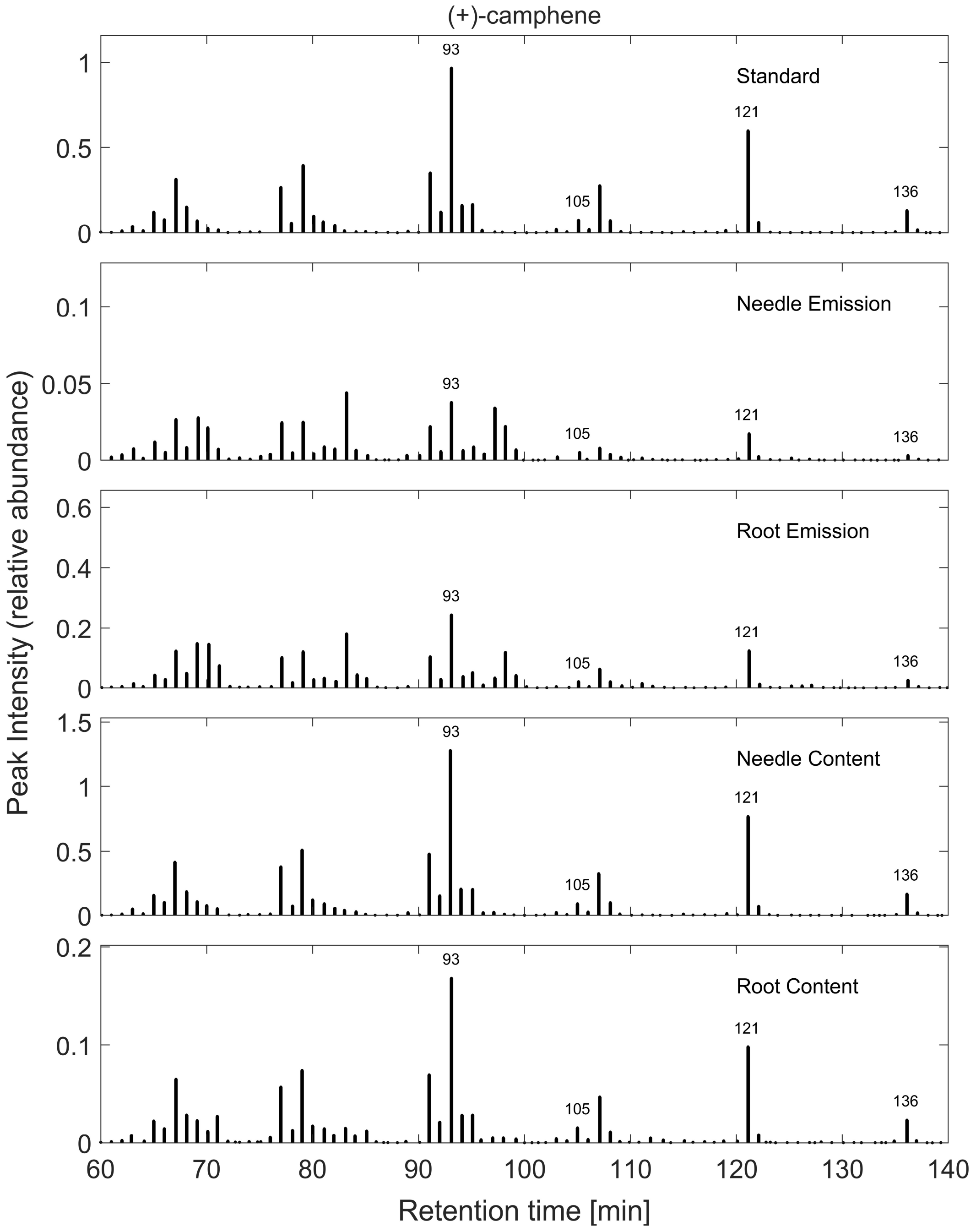


**Fig. S8** GCMS spectrum of (+)-camphene from different samples of *Picea abies* and a Standard. Peak intensity is displayed in $10^5$-steps.

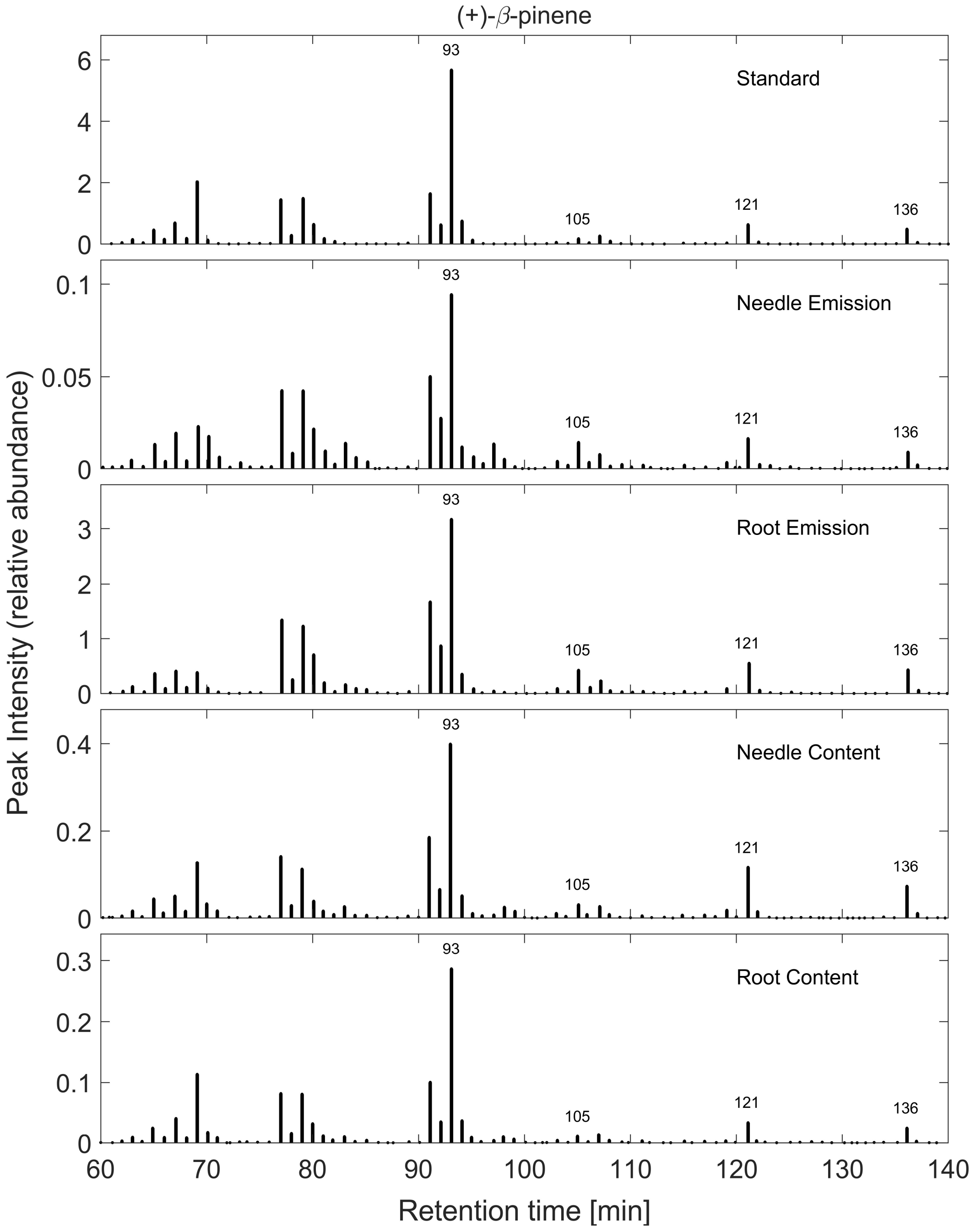


**Fig. S9** GCMS spectrum of (+)-β-pinene from different samples of *Picea abies* and a Standard. Peak intensity is displayed in $10^5$-steps.

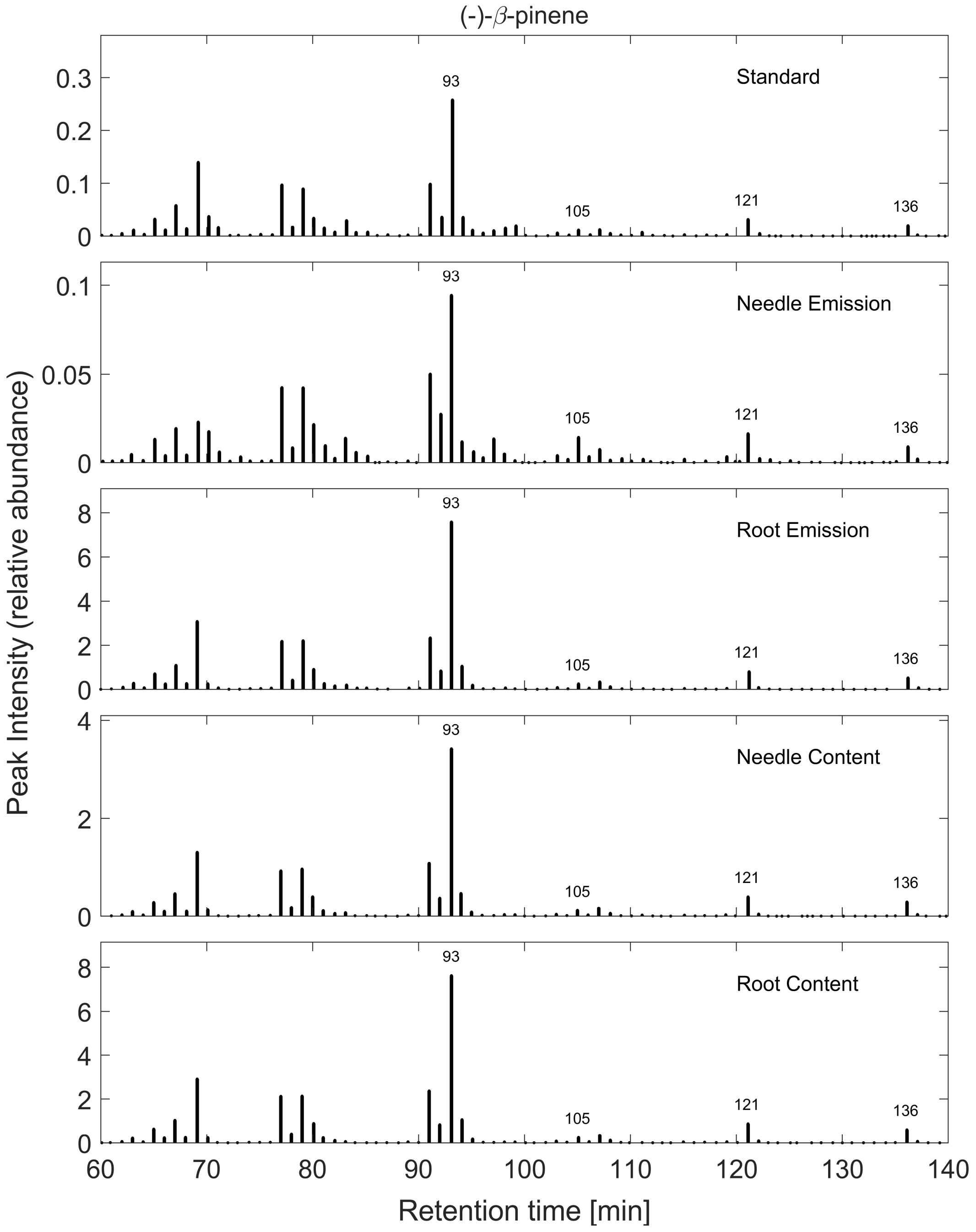


**Fig. S10** GCMS spectrum of (-)-β-pinene from different samples of *Picea abies* and a Standard. Peak intensity is displayed in $10^5$-steps.

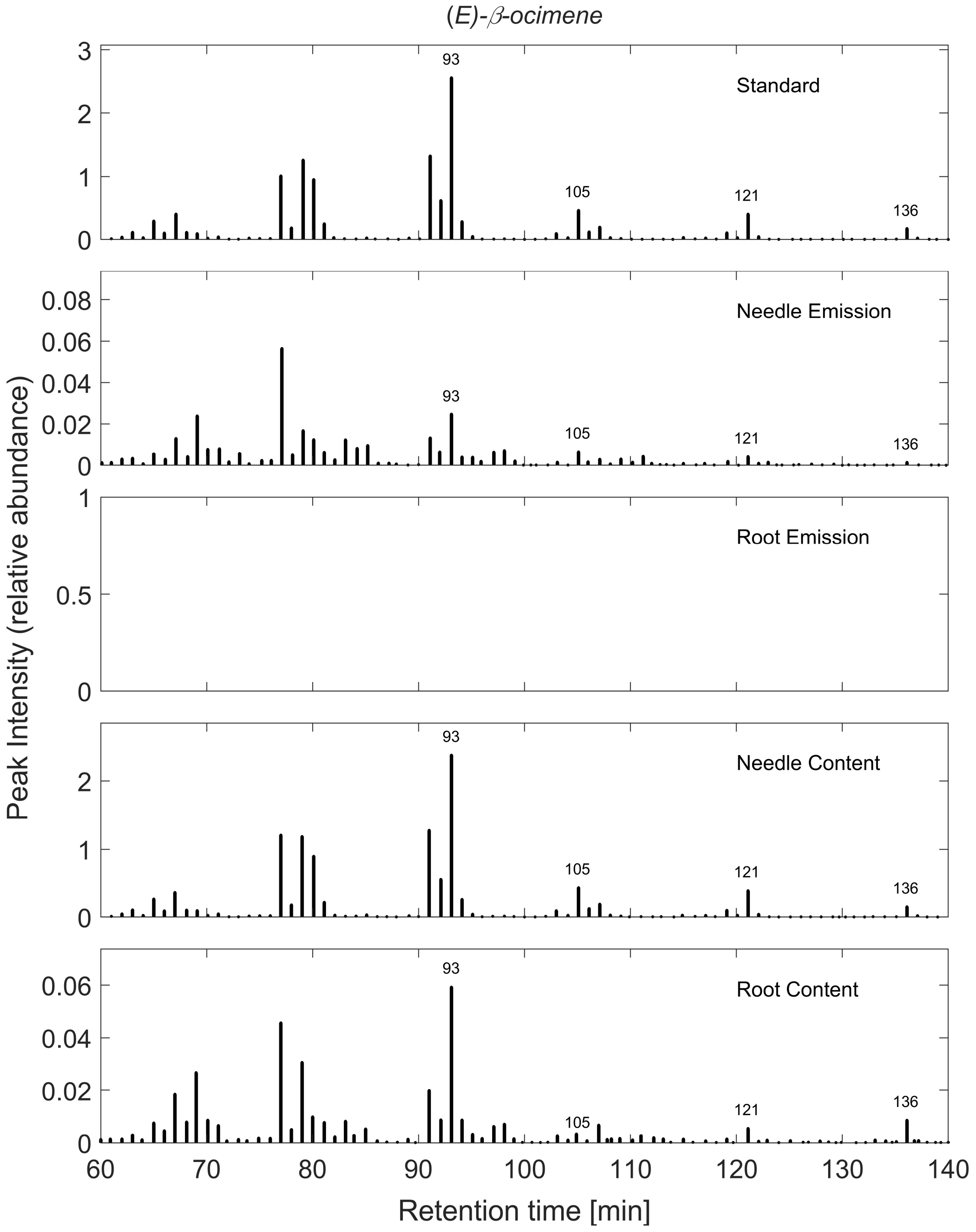


**Fig. S11** GCMS spectrum of (*E*)-β-ocimene from different samples of *Picea abies* and a Standard. Peak intensity is displayed in $10^5$-steps. Empty graphs indicate that the terpene was not identified in the sample.

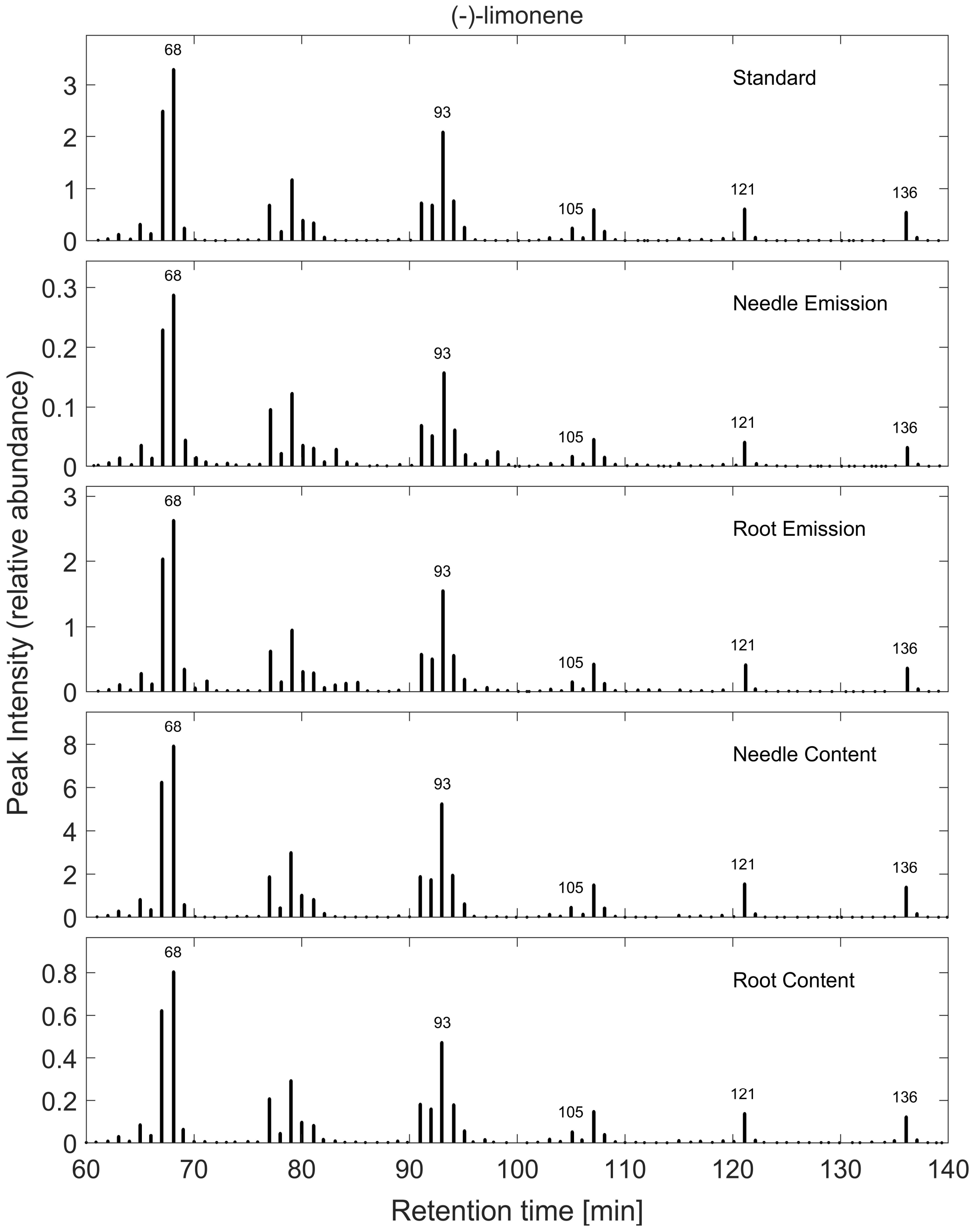


**Fig. S12** GCMS spectrum of (-)-limonene from different samples of *Picea abies* and a Standard. Peak intensity is displayed in $10^5$-steps.

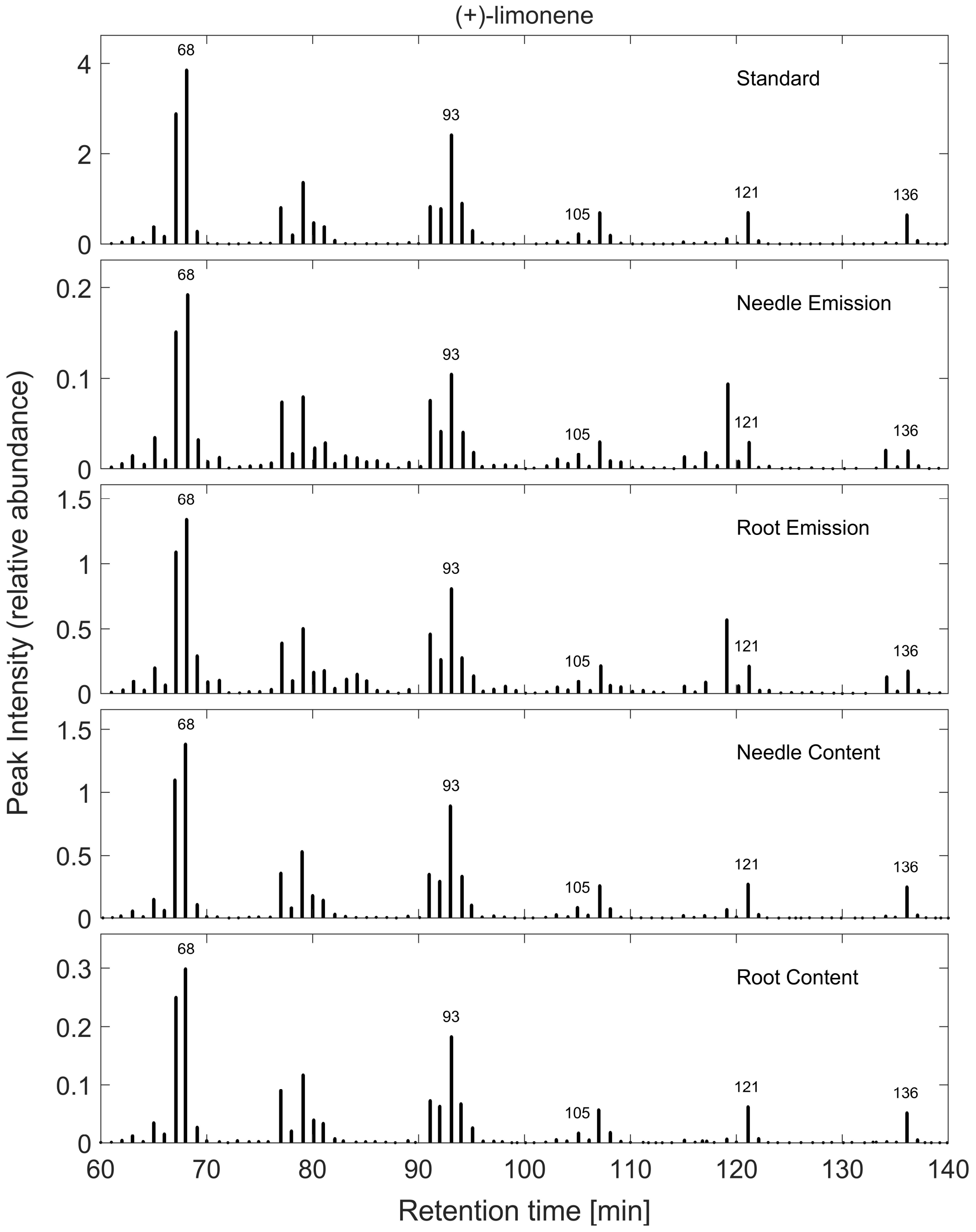


**Fig. S13** GCMS spectrum of (+)-limonene from different samples of *Picea abies* and a Standard. Peak intensity is displayed in $10^5$-steps.

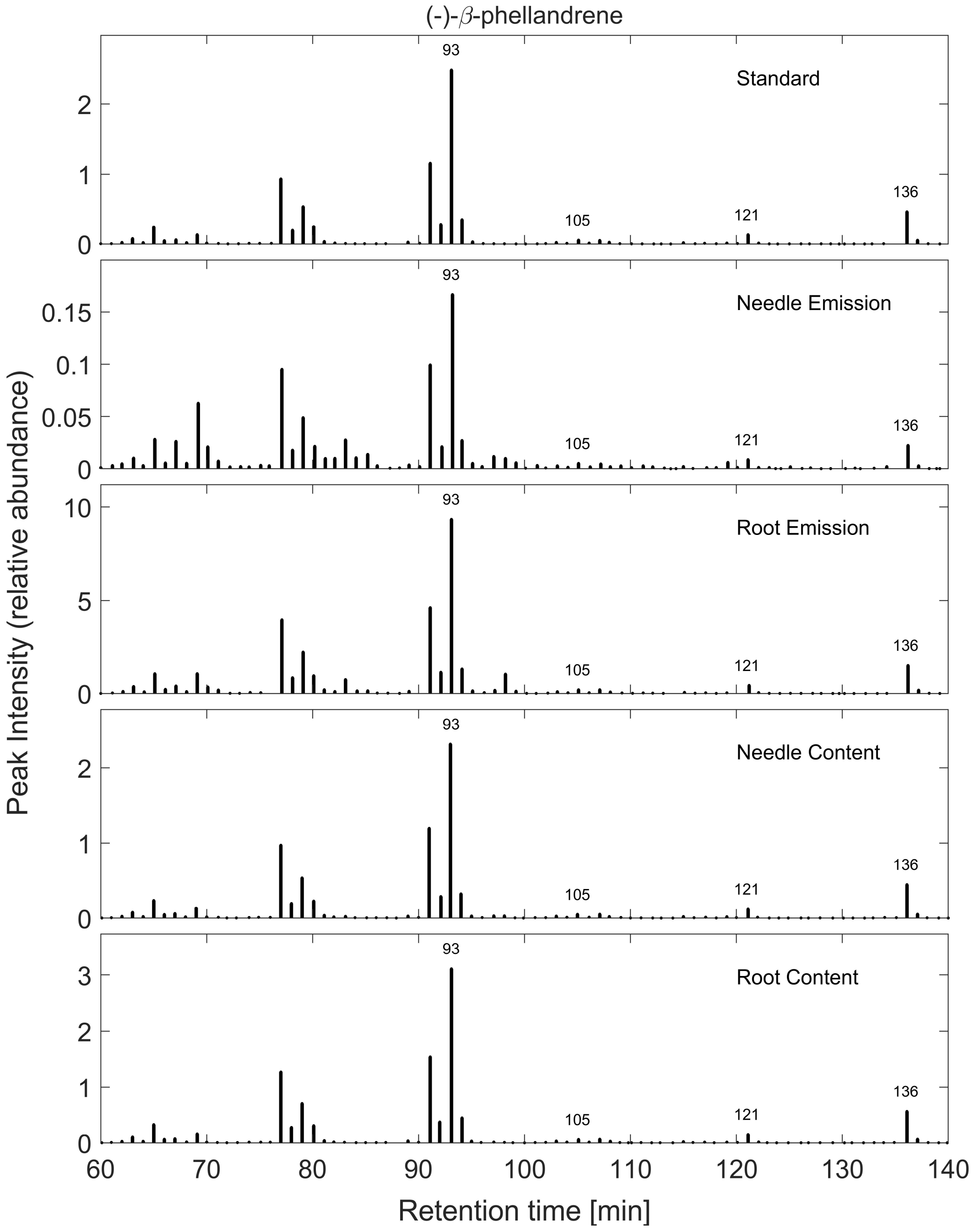


**Fig. S14** GCMS spectrum of (-)-β-phellandrene from different samples of *Picea abies* and a Standard. Peak intensity is displayed in $10^5$-steps.

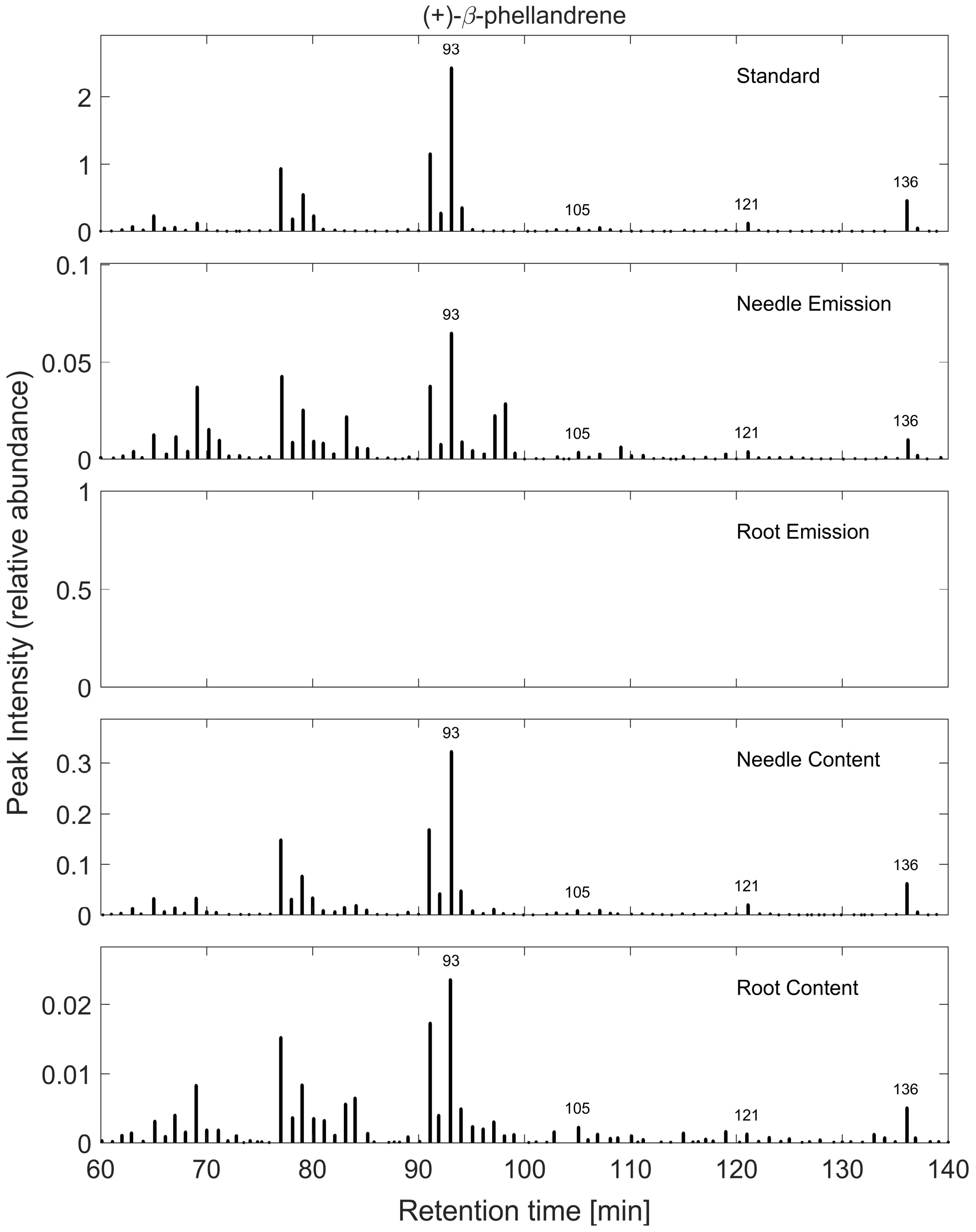


**Fig. S15** GCMS spectrum of (+)-β-phellandrene from different samples of *Picea abies* and a Standard. Peak intensity is displayed in $10^5$-steps. Empty graphs indicate that the terpene was not identified in the sample.

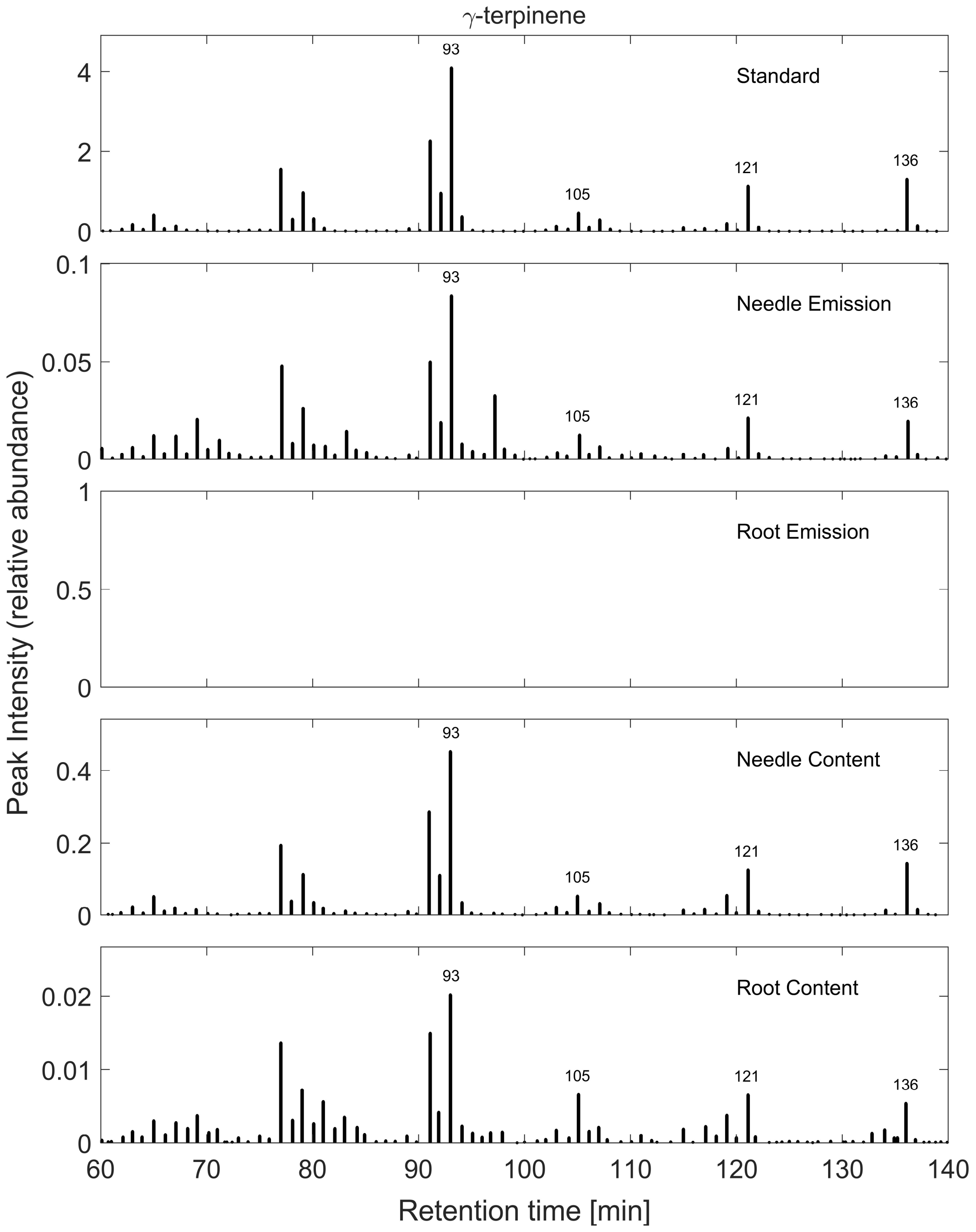


**Fig. S16** GCMS spectrum of γ-terpinene from different samples of *Picea abies* and a Standard. Peak intensity is displayed in $10^5$-steps. Empty graphs indicate that the terpene was not identified in the sample.
